# Growth costs of protein allocation in models of growing cells

**DOI:** 10.64898/2026.04.21.719900

**Authors:** Diana Széliová, Wolfram Liebermeister, Martin J. Lercher, Hugo Dourado

## Abstract

Microbial growth depends on how cells allocate their limited proteome among competing functions. Although many proteins are expressed near levels that maximize growth, deviations from these optima are common and can impose substantial fitness costs. Modeling the growth costs of suboptimal protein allocation remains challenging because protein expression influences growth through multiple interacting mechanisms, including biosynthetic demands, enzyme kinetics, and limits on cellular density. Here we analyze these effects using growth balance analysis (GBA), a nonlinear framework that predicts steady-state growth and biomass composition of coarse-grained microbial models from basic cellular constraints. The optimal biomass compositions predicted for these models include not only protein concentrations but also the reactant concentrations required to saturate enzymes through nonlinear rate laws, thereby capturing more complex resource trade-offs than in conventional linear resource allocation models. Using these GBA models, we compute optimal biomass compositions with individual proteins fixed at suboptimal levels. The resulting relationships between suboptimal protein allocation and growth rate are consistent with qualitative experimental patterns in bacteria, with growth effects depending on protein function (including idle proteins), environmental conditions, toxic byproducts, and alternative reactions. These results indicate a practical and tractable approach for modeling growth costs arising from suboptimal protein allocation, and suggest a basis for predictive modeling in metabolic engineering and synthetic biology.

## Introduction

Proteins play a central role in determining microbial growth, in particular by composing the catalysts of biochemical reactions such as the transporters exchanging resources with the environment, internal enzymatic reactions involved in metabolism and biosynthesis, and the ribosome which is itself involved in synthesizing proteins (1, 2). Proteins are also themselves costly components for the cells, since their synthesis requires products from metabolism and biosynthesis (such as ATP and precursors) which could otherwise go into the synthesis of other key macromolecules such as RNA and DNA (1, 3). Microbes face constant selective pressures on how to balance such resource allocation among competing cellular functions (2, 4, 5). This trade-off has been interpreted in theoretical studies as a growth optimization problem under physico-chemical constraints (1, 3, 6–10). Experimental studies in bacteria and yeast have shown that many proteins are indeed expressed near levels that maximize their growth rate in a given environment, since forcing overexpression or underexpression typically reduces growth rate when compared to wild types (2, 11–14). However, it has also been observed that microbes frequently express proteins that appear unnecessary under the experimental conditions, such as catabolic proteins and transcription factors associated with alternative nutrient sources, or stress response proteins (15– 17). A theoretical understanding of how such deviations from optimal protein expression affect growth quantitatively remains an important problem in microbial physiology and metabolic engineering, and could potentially guide strategies to improve metabolic efficiency (18).

Changes in protein expression affect cell growth through multiple interacting mechanisms (19). These include the direct increase in the resources required for transcription and translation, which depend primarily on protein size and composition (20) and can in principle be estimated from precursor and energy requirements or measured in cell-free systems (21, 22). However, additional costs associated with protein folding and degradation remain difficult to quantify (23). In addition to these direct biosynthetic costs, indirect costs arise from system-level effects. Increasing the concentration of one protein limits the concentrations of others through molecular crowding, and it can alter reaction fluxes, inhibit cellular functions, or perturb membranes (14, 24, 25). Over-expression of some enzymes also imposes metabolic burdens through the accumulation or depletion of metabolites that reduce growth rate (25). These include engineered products such as biofuels (24) and native metabolites such as methyl-glyoxal at elevated levels (26), which damage cellular components and impair growth. Consequently, overexpression of different proteins affects growth rate to varying degrees (21, 23–25, 27, 28). Understanding these complex interactions between protein allocation and growth remains challenging. Some studies provide a local understanding of these phenomena through mathematical models and experiments on the suboptimal expression of individual proteins (12), but they do not explain how global proteome allocation strategies emerge.

Mathematical models of cellular reaction networks provide a framework for illuminating the connection between growth rate and proteome allocation at the level of the whole cell. Some constraint-based metabolic models build on classical flux balance analysis (FBA) (29) by incorporating linear constraints that relate proteome allocation to growth under the assumption of a predefined biomass composition (6–9). Resource balance analysis (RBA) further incorporates ribosomal protein synthesis costs and captures resource allocation in a self-consistent manner for macromolecules including proteins, RNA, and DNA (3). These frameworks have provided important insights into growth laws and proteome organization through linear optimization procedures that remain computationally feasible for large and detailed models. These insights include, for example, predictions of optimal proteome allocation strategies, metabolic trade-offs, and the impact of resource limitations on cellular physiology (3, 6– 9). However, important aspects of the relationship between proteome allocation and growth remain unexplained for two main reasons related to the intrinsic nonlinear nature of the problem. First, from a practical point of view, many of the physical parameters required to construct detailed nonlinear cell models are not known from the literature, constituting a major open problem in metabolic modeling (30). Second, from a theoretical point of view, the exact mathematical relationship between proteome allocation and growth rate has only recently been identified (31).

Previous theoretical studies have treated the explicit mathematical relationship between proteome allocation and cell growth rate in different ways. For example, studies on protein expression noise typically assume the existence of an unknown growth-rate function *µ*(***ϕ***) depending on the proteome mass fractions ***ϕ*** allocated to different reactions, and approximate this function locally as linear around the mean protein expression levels, so that proportional changes in protein concentrations produce proportional changes in growth rate (32). Molenaar et al. (1) previously introduced a general numerical approach to solve this problem without such assumptions for small coarse-grained self-replicator models, showing that major economic patterns of cellular resource allocation can be captured from first principles. We later extended this approach by deriving an exact analytical solution for the relationship between growth rate and entire cell state, including proteome allocation, in the same general class of models through the growth balance analysis (GBA) framework (10), based on the first principles of balanced growth optimization under the constraints of mass conservation, nonlinear kinetic rate laws, and fixed cell density.

The key mathematical advance in GBA is the explicit formulation of the growth-rate function *µ*(**q**) in terms of independent state variables **q**, which enables analytical and numerical studies of growth-rate sensitivity to changes in cellular state in a general setting (see Methods for mathematical details). We have previously used this framework to characterize the analytical properties of optimal growth states and, using small coarse-grained GBA models whose few parameters are available in the literature, to predict ribosome concentrations in microbes across different growth rates with good agreement with experimental data (10).

An optimization of cellular states can also be formulated based on elementary growth modes (EGM), introduced by de Groot et al. (33) for kinetic models. A variant of EGMs has also been developed for linear cell models (34).

Here, we apply the GBA framework beyond optimal balanced-growth states to characterize suboptimal microbial states arising from protein under- or overexpression. Specifically, we constrain the mass fraction of a selected protein to a suboptimal value while optimizing the remaining proteome allocation, allowing us to quantify the resulting growth-rate response. This broadens previous GBA formulations, which were limited to optimal states, by enabling systematic analysis of suboptimal states in mechanistic models of growing cells.

## Results

We investigate the growth costs associated with deviations from optimal protein expression using a set of minimal models of microbial cells. We examine both under- and over-expression of active enzymes, as well as the expression of idle proteins, and quantify how the resulting growth costs depend on environmental and network features. We study how these costs vary with substrate availability, the presence of alternative reactions, and trade-offs arising from the allocation of expression between two proteins.

Throughout the following analysis, the term “proteins” is used broadly to denote active enzymes, molecular machines, and idle proteins. When enzymes contain additional components such as RNA (for example the ribosome), our analysis refers specifically to the protein component.

### Nonlinear resource allocation model of a microbial cell

We investigated the growth costs of protein allocation by first constructing a GBA model that contains basic components of a generic microbial cell (referred to as “base model”), including basic metabolism and gene expression processes (Figure 1). In this model, the cell imports nutrients S_ext_ via a transporter, TS. The intracellular “lumped metabolite” S represents a total pool of all cell compounds that are not explicitly modeled. The ATPS reaction converts S into a waste product W, which is excreted, and generates energy in the form of ATP from ADP. The AAS reaction synthesizes amino acids (AA) from S. The NTS reaction, powered by ATP, produces nucleotides (NT), and ADPS synthesizes ADP from NT. Nucleotides are used by RNA polymerase (RNAP) to synthesize RNA and by DNA polymerase (DNAP) to synthesize DNA. Finally, ribosomes (reaction R) use amino acids and ATP to produce the proteome P. This reaction represents the total synthesis of all proteins in the model, and P denotes the total modeled proteome (a sum of all proteins). In this model, we implicitly assume that there is another, constant proteome fraction (often called the Q sector, “other”, or housekeeping proteins), which constitutes about half of the cellular proteome (2), but is not explicitly included in our formulas. Unless stated otherwise, “proteome” refers to the modeled, growth-associated proteome fraction.

**Fig. 1.**
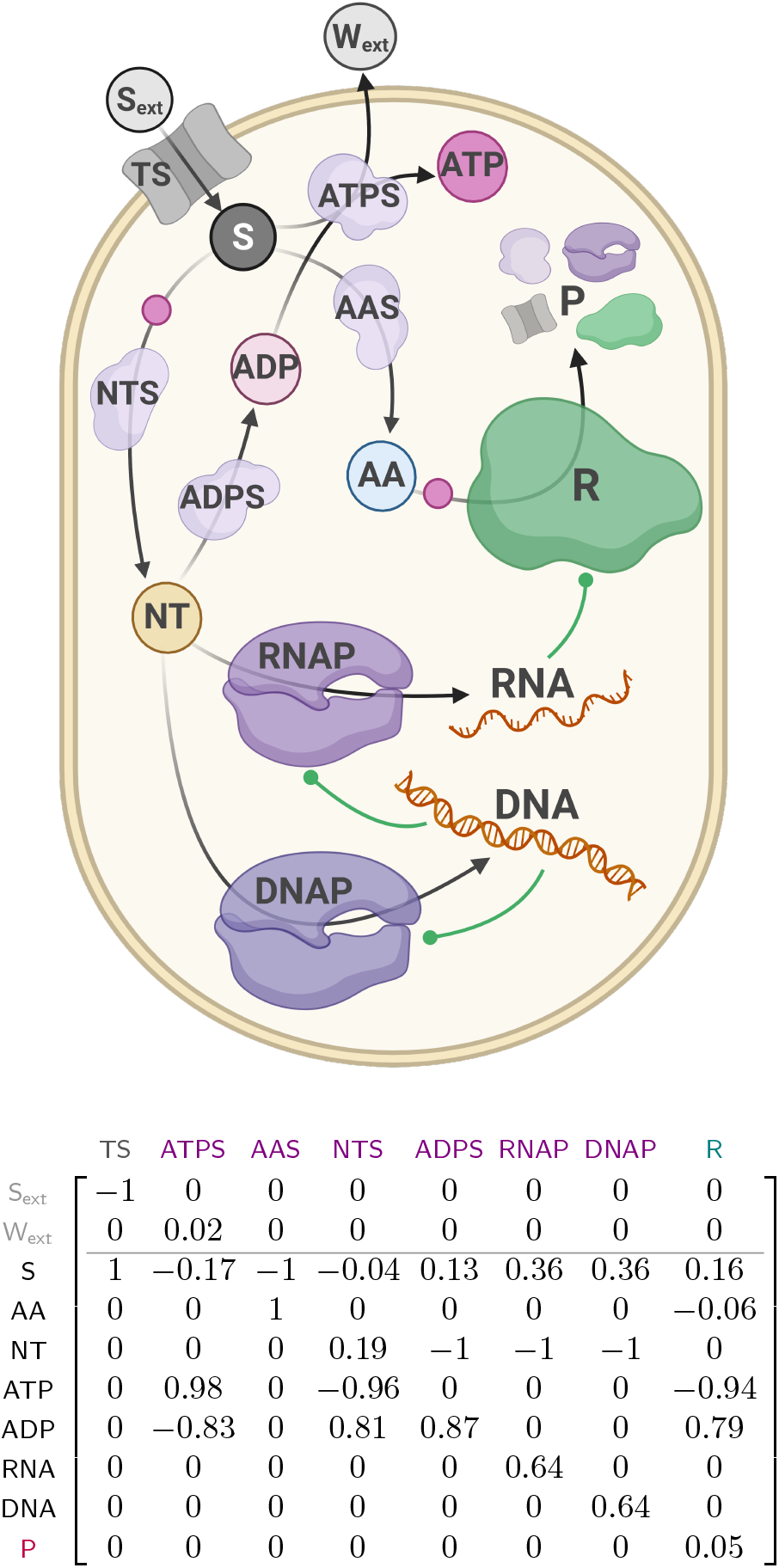
A base model of a self-replicating cell, and its corresponding total mass fraction matrix M_total_. The reactions between metabolites (circles) are catalyzed by specific catalytic proteins. Each reaction has an explicit kinetic rate law that links its flux to the metabolite concentrations and to the abundance of its catalyzing protein. The cell takes up small molecules S_ext_ via a transporter (TS), and converts them into amino acids (AA) and nucleotides (NT) through amino acid synthesis (AAS) and nucleotide synthesis (NTS) reactions. The internal species S is a lumped variable that represents several different internal metabolites. Nucleotides are converted into ADP by the ADP synthesis reaction (ADPS). The ATP synthesis reaction (ATPS) uses S and ADP to generate ATP, producing a waste product W_ext_ that is immediately excreted. Nucleotides are then used by DNA polymerase (DNAP) and RNA polymerase (RNAP) to synthesize DNA and RNA, respectively. Amino acids and ATP are used by ribosomes (R) to synthesize the proteome (P), which represents the total modeled proteome, defined as the sum of all proteins in the model. Small pink circles on top of the arrows indicate that a reaction requires ATP. Green lines with circles represent that a molecule is required for a reaction to proceed, but is not consumed in the reaction – DNA is required by both polymerases as a template and RNA is required as a component of the ribosome. In GBA, the growth rate is maximized by optimizing the reaction fluxes under the constraints of balanced growth, constant cell density, and the kinetic rate laws. See SI for details about constructing the **M**_total_ matrix. The density and kinetic parameters are based on *E. coli* data (see Methods and Tables 2 and 3). Cell figure created with biorender.com

Fig. 1 shows the mass fraction matrix of the base model. The full model description is given in the Methods section.

It contains all constraints, kinetic rate laws, and parameter values (Tables 2 and 3).

**Table 1.** Overview of mathematical symbols.

| Symbol | Description [units] |
| --- | --- |
| $\mathbf{a}$ | external concentration vector [g/L] |
| $\mathbf{b}$ | biomass fraction vector |
| $\mathbf{c}$ | metabolite concentration vector [g/L] |
| $e_{\text{sat}}$ | enzyme saturation |
| $k_{\text{cat}}$ | turnover number 1/h |
| $\mathbf{K}$ | Michaelis constant matrix [g/L] |
| $\mathbf{A}$ | activation constant matrix [g/L] |
| $\mathbf{I}$ | inhibition constant matrix [g/L] |
| $\mathbf{M}_{\text{total}}$ | total mass fraction matrix |
| $\mathbf{M}$ | mass fraction matrix for internal reactants |
| $\mathbf{p}$ | protein concentrations vector [g/L] |
| $\mathbf{q}$ | flux fraction vector |
| $r_{\text{sat}}$ | ratio of metabolite concentration to $K$ |
| $\mathbf{s}$ | vector with column sums of $\mathbf{M}$ |
| $\mathbf{v}$ | flux vector [g/(Lh)] |
| $\mu$ | growth rate [1/h] |
| $\rho$ | mass density [g/L] |
| $\boldsymbol{\tau}$ | turnover time vector [h] |
| $\boldsymbol{\phi}$ | proteome mass fraction vector |
| Index | Description |
| $i$ | internal metabolites |
| $j$ | external metabolites |
| $l$ | reactions |
| $P$ | proteome |
| $R$ | ribosome reaction |

**Table 2.**
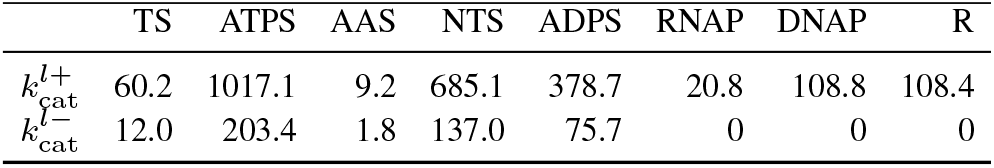
Estimated turnover numbers in g product/(g protein h).

|  | TS | ATPS | AAS | NTS | ADPS | RNAP | DNAP | R |
| --- | --- | --- | --- | --- | --- | --- | --- | --- |
| $k_{cat}^{l+}$ | 60.2 | 1017.1 | 9.2 | 685.1 | 378.7 | 20.8 | 108.8 | 108.4 |
| $k_{cat}^{l-}$ | 12.0 | 203.4 | 1.8 | 137.0 | 75.7 | 0 | 0 | 0 |

**Table 3.**
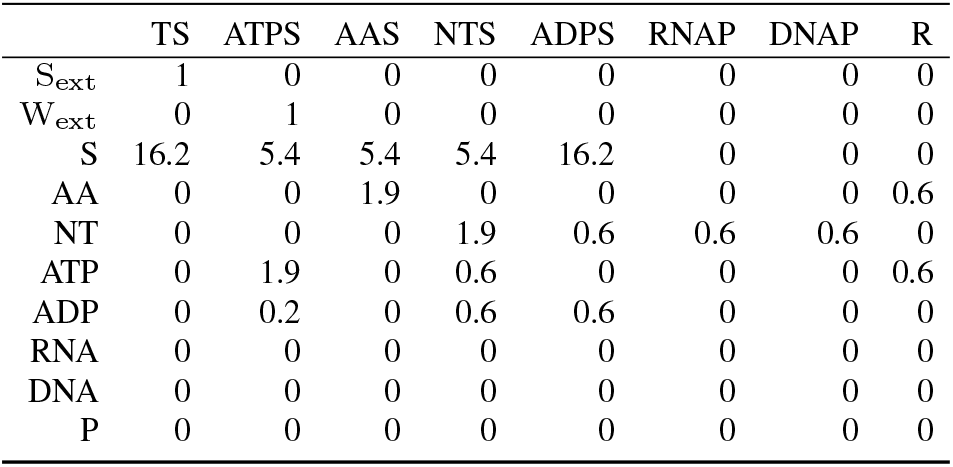
Estimated K values assuming reversible kinetics in g/L.

In the main text, we assume that all metabolic reactions (TS, AAS, NTS, ADPS, and ATPS) follow reversible kinetics (Eq. (13)), meaning that the reaction rate is reduced by the accumulation of the product. In the SI, we also show results with irreversible kinetics where no product inhibition is included. The macromolecular reactions DNAP, RNAP, and R are always treated as irreversible. DNA is essential as a template for its own replication and for transcription. To represent this in the model, a DNA-dependent regulatory term appears in the rate laws for DNAP and RNAP (Eq. (13)) to reflect its requirement for these processes. In GBA, ribosomes are represented solely as proteins. To account for their rRNA content and the requirement of mRNA and tRNA for function, we incorporated an RNA-dependent term into the ribosome rate law (reaction R, Eq. (13)). This term is a pragmatic way to link ribosome function to the availability of all necessary RNA species, capturing overall resource demands without modeling ribosome assembly or translation in detail. Although the model represents a generic microbial cell, we applied it to *Escherichia coli*, estimating the kinetic parameters of its reactions based on *E. coli* data, including proteomics, biomass composition, and known turnover numbers (see Methods Section Parameters and experimental data, and Supplementary Information).

To determine whether the model reproduces well-known experimental relationships, we examined the fraction of growth-associated protein mass allocated to ribosomal proteins, and how the RNA/protein ratio depends on the growth rate (which is set by a fixed extracellular concentration of the carbon source S_ext_). Both quantities vary linearly with the growth rate (Figure S4), consistent with experimental data (35–37). The model also captures the linear increase in the ribosomal proteome fraction in the presence of a translation inhibitor (Figure S5) (2). The kinetic parameters were calibrated to the growth rate and biomass composition of *E. coli* (see Methods). The dependence of the ribosomal fraction and the RNA/protein ratio on growth rate, and the response to a translation inhibitor, were not used in this calibration. Together, these results show that the model reproduces these growth laws without explicit implementation of the laws as model constraints.

### The nonlinear growth cost of suboptimal protein expression

To compute the growth cost of individual proteins, we systematically varied, for each protein, the mass fraction of the modeled proteome allocated to this protein (referred to below as the proteome fraction or protein expression level), and calculated the maximal growth rate achievable through an optimal allocation of the remaining cellular mass density to all other cellular components (Figure 2). For every protein, the curve has a maximum point corresponding to the optimal expression that GBA would predict if no suboptimal proteome fractions were enforced.

**Fig. 2.**
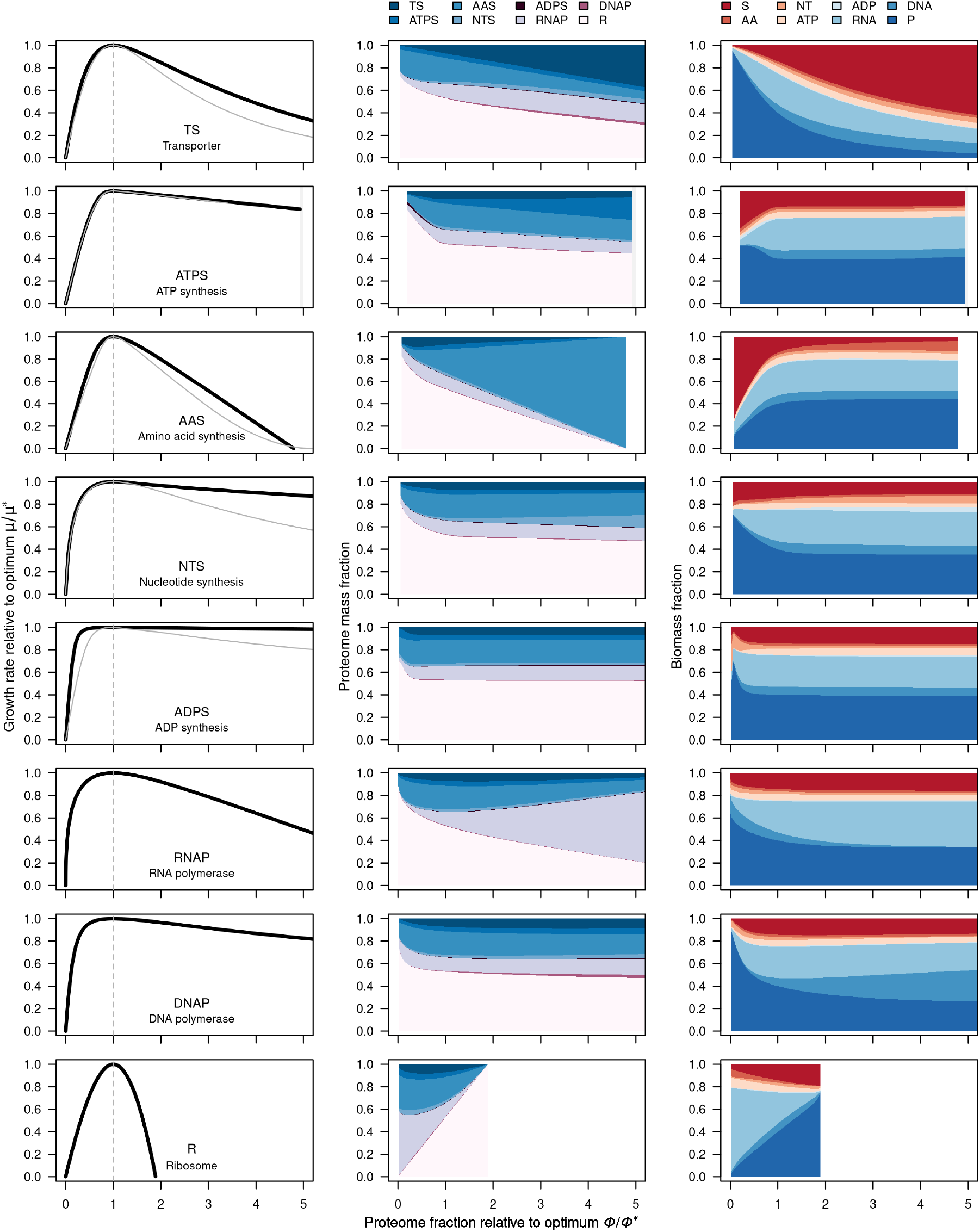
The growth cost of suboptimal protein expression, emerging from calculations based on first principles. The first column shows, for each reaction, the normalized growth rate (normalized to maximum growth rate) as a function of *ϕ* relative to the optimal *ϕ*^∗^. For each point, the proteome fraction *ϕ* of the plotted protein is fixed, and the fractions of all other proteins are re-optimized to maximize growth rate. The black curves are simulations with reversible kinetics and the gray curves with irreversible kinetics for comparison. Full results with irreversible kinetics are shown in Figures S7 and S9. The second column shows the proteome composition, and the third column shows the biomass composition, where P represents the mass fraction of the proteome in the biomass. Note that gaps or noise can occur when the calculation does not converge – the further *ϕ* is from the optimum, the more likely the optimization solver is to fail to find a solution. For visualization purposes, growth rates were extrapolated to zero and data points were connected. Results with individual data points over the full range of *ϕ* values are provided in the supplementary material (Figure S8). Numerical results for ATPS were not found for higher proteome fractions, and ribosome overexpression is limited before it doubles since it completely dominates the proteome already below this point.

The resulting protein-cost curves are nonlinear, and their overall shape matches experimental observations (11, 27), see Figure S6. Notably, this characteristic shape already emerges in a minimal toy model containing only three reactions (Figure S2). We also compared GBA with RBA (as formulated in (38)), enzyme-constrained FBA, and the kinetic COBRA-k framework (39) on this toy model (Supplementary Section “Comparison with other modeling approaches”). The linear methods give two straight segments with a sharp optimum, and the kinetic COBRA-k model reproduces the rounded shape of GBA around the optimum only partly.

As a protein’s proteome fraction *ϕ* approaches zero, the growth rate sharply declines to zero because each protein has a functional role in the model and is therefore essential – the growth rate reduction results from a declining benefit. In later sections, we investigate how introducing alternative enzymes with distinct kinetic parameters modifies this behavior.

For comparison, we used both reversible and irreversible rate laws to test whether the resulting growth costs differ substantially. In the first column of Figure 2, we observed that reversibility had little effect near the optimum, but that irreversible kinetics resulted in higher growth costs at greater protein overexpression. For reversible kinetics (black curves), growth rate declines approximately linearly (see Figure S8 for results across the full *ϕ* range); for irreversible kinetics, the decline is nonlinear for several metabolic reactions (TS, AAS, and NTS; see gray curves in Figure 2).

Functional proteins can impose indirect costs on the cell beyond their biosynthetic cost, for example by perturbing metabolic processes (referred to as metabolic burden) (25). In our framework, these effects arise naturally from the nonlinear resource-allocation model through changes in biomass allocation and reaction kinetics, including inhibition due to metabolite accumulation or regulatory mechanisms.

To understand the origin of these differences between reversible and irreversible kinetics, we examined how biomass composition changes with protein expression levels. With reversible rate laws, metabolite accumulation remained limited across the range of protein expression levels (Figures 2 and S8), whereas irreversible rate laws can lead to much larger increases in metabolite mass fractions, with the general pool S or amino acids AA dominating the biomass in some cases (Supplementary Figures S7 and S9). This difference arises because, in reversible rate laws, product accumulation reduces the reaction rate, providing a buffering mechanism that limits excessive metabolite accumulation.

These model predictions are consistent with experimental observations: systematic overexpression of glycolytic enzymes in yeast resulted in only minor changes in metabolite concentrations, which was hypothesized to arise from reversible kinetics (14). In line with this, we also expect that models with reversible kinetics would in general maintain a more stable total protein content under perturbations in enzymatic expression, due to this distributed response across the reaction network under a localized perturbation (see Figure 2 for an illustrative example). This stable total protein content following protein overexpression has been reported in yeast (40). Despite their potential to capture such buffering effects more realistically, reversible kinetic models require substantially more parameters, which are generally not available from the literature.

### Growth costs depend on environmental conditions

Protein expression costs and benefits depend strongly on a cell’s environment: proteins that are essential in one condition may be unnecessary or harmful in others. Dekel and Alon (12) demonstrated this by studying the lactose utilization system, a set of proteins for lactose uptake and its regulation, encoded on a single operon. In their experiments, cells were grown on glycerol as the primary carbon source, and the *lac* operon was overexpressed either with or without lactose added to the growth medium. It turned out that expressing the *lac* system reduced growth in the absence of lactose, but increased the growth rate when lactose was present.

To model a corresponding scenario, we introduced an alternative carbon source, L_ext_, into the base model. For simplicity, transport and catabolism of L_ext_ are described by a single reaction catalyzed by a single effective protein (“LAC protein”):

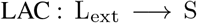

Glycerol utilization is represented by the reaction already in the model (S_ext_ → S).

Figure 3a shows the predicted growth rate as a function of *ϕ*_LAC_ for different L_ext_ concentrations. At low L_ext_ (much below the transporter’s Michaelis constant 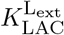), expression of LAC incurs a nearly linear growth cost, consistent with observations from LacZ protein overexpression experiments (2). At high L_ext_, the benefit initially increases steeply, then plateaus and eventually declines at high *ϕ*_LAC_, reflecting a cost–benefit trade-off. The optimal *ϕ*_LAC_ shifts to lower values as L_ext_ increases: when the substrate saturates the transporter, even a small allocation to LAC provides sufficient carbon flux. Beyond this optimum, costs outweigh benefits, and growth rate drops below the baseline.

**Fig. 3.**
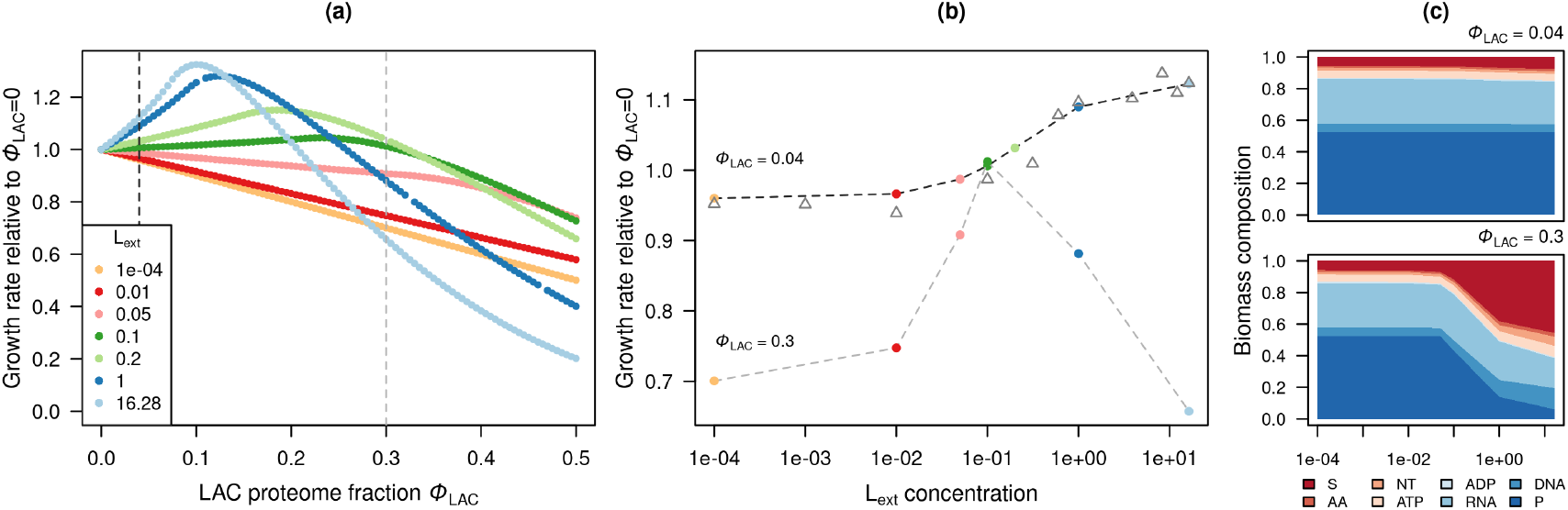
The growth cost of lactose utilization (LAC) proteins depends on environmental conditions. **(a)** Relative growth rate (normalized to the growth rate when *ϕ*_LAC_ = 0) as a function of proteome allocation to LAC (*ϕ*_LAC_) at different L_ext_ concentrations. Higher L_ext_ levels shift the optimum toward lower *ϕ*_LAC_. **(b)** Relative growth rate as a function of L_ext_ concentration. At *ϕ*_LAC_ = 0.04, the curve reproduces experimental results from (12) shown as gray triangles. (Note that experimentally, LAC protein overexpression was induced using saturating concentrations of IPTG, and the corresponding proteome fraction was not measured.) **(c)** Biomass composition for each *ϕ* shows metabolic perturbation at high *ϕ*_LAC_.

To mirror the analysis by Dekel and Alon (12), we plotted the predicted growth rates in normalized form (Figure 3 (b)), relative to the growth rate in the absence of LAC proteins (*ϕ*_LAC_ = 0) for *ϕ*_LAC_ = 0.04 and 0.3. In our model, the glycerol transporter was assigned kinetic parameters that reproduce the experimentally observed growth rate relative to *ϕ*_LAC_ = 0 at maximal L_ext_ concentration (see Methods). In Ref. (12), full *lac* operon induction caused a 4.5% reduction in growth rate. Because this work did not measure the *lac* proteome fraction, we estimated it using data from (2). In that dataset, a 4.5% reduction in growth rate corresponds to roughly 2% of the total proteome being allocated to *β*-galactosidase. Excluding the constant housekeeping fraction (Q), which is not included in this model, this corresponds to about 4% of the modeled growth associated proteome.

At *ϕ*_LAC_ = 4%, the model reproduces the findings of Dekel and Alon (12): at low L_ext_, the enzyme is purely burdensome, while increasing L_ext_ makes it beneficial for growth by enabling lactose utilization. As an additional comparison, we considered the extreme case at *ϕ*_LAC_ = 0.3, where the metabolic burden exceeds the baseline cost at L_ext_ = 10^−4^. This value represents approximately the maximum proteome fraction observed for overexpressed *β*-galactosidase experimentally (2). However, we note that such high expression would not be expected for the full *lac* operon, which also requires transporter expression and is therefore constrained by membrane availability.

The biomass composition plots in Figure 3 (c) show accumulation of intracellular S at high *ϕ*_LAC_, explaining why growth rates decline faster under high-lactose conditions. However, such a high accumulation of an intermediate (up to almost half of the biomass) is unrealistic, as cells would be expected to limit this accumulation through regulatory mechanisms or by exporting excess substrate. Introducing an export transporter into our model reduces the intracellular accumulation of S and alleviates the associated growth cost (Figure S11).

The model structure used in this section can represent the uptake of two different carbon sources at different concentrations. In the current model, both carbon sources produce the same intracellular metabolite, so the two carbon sources are metabolized through the same downstream reactions. Alternative reactions could be introduced to represent distinct metabolic pathways.

### Idle protein expression results in a linear growth cost

In some bioengineering applications, proteins are expressed that do not perform any native function in the host cell. Examples include marker proteins such as green fluorescent protein (GFP) and biotechnologically relevant products such as antibodies.

To simulate this scenario, we introduced a nonfunctional protein into the model. To enable direct comparison with experimental data, we also included the non-growth-associated proteome (Q sector) in the model, and fixed its abundance to 55% of the total proteome. The chain length of the idle protein does not affect the model predictions, because GBA describes proteins through their mass fractions and assumes that ATP and amino acid costs scale approximately with protein mass.

Figure 4a shows that increasing the proteome fraction allocated to the idle protein causes an approximately linear decrease in growth rate. Notably, the biomass composition remains essentially unchanged, including the total protein fraction (Figure 4c). This agrees with experimental observations showing that overexpression of idle proteins leads to proportional growth reductions (2, 5, 19), while causing only minor changes in the protein, RNA, and DNA fractions of the biomass (40, 41).

**Fig. 4.**
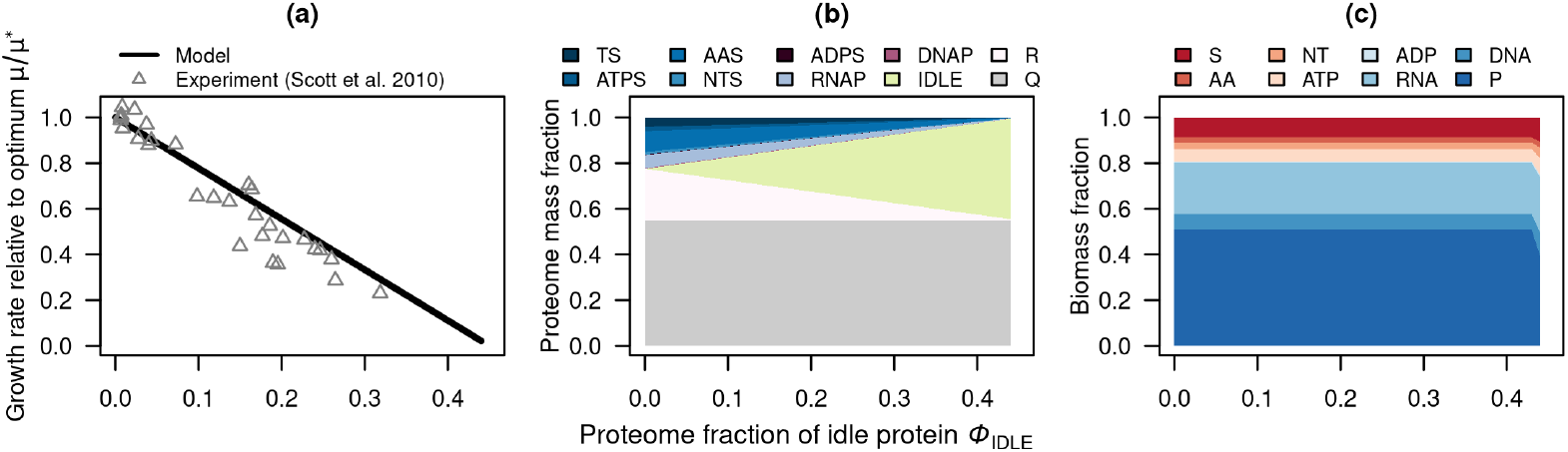
Expression of an idle protein. Increasing the proteome fraction of an unnecessary protein leads to an almost linear decrease in growth rate (a), consistent with experimental data (2). Each gray triangle in (a) is one measurement from Scott et al. (2010) (2) of the growth rate at a given proteome fraction of the overexpressed protein (*β*-galactosidase). The data cover different growth conditions, and each growth rate is normalized to the maximum growth rate of its condition, measured without overexpression. This simple behavior results from the protein occupying an increasing fraction of the proteome (b), while the remaining biomass composition, and therefore the catalytic efficiencies of all proteins, remains unchanged (c).

Because the cost of idle protein expression is linear, the slope of the growth-rate reduction depends on the fraction of the proteome that remains available for growth-associated functions. Assuming a constant non-growth-associated (*Q*) proteome fraction of 55%, consistent with proteomic estimates, the model predictions in Figure 4a quantitatively reproduce the growth costs reported in (2). In contrast, without a Q sector, the growth rate decreases linearly to zero as the idle protein occupies the entire proteome (Figure S12).

These results indicate that idle protein expression primarily reduces the proteome fraction available for functional proteins, without substantially perturbing the allocation of the remaining biomass. Consequently, enzyme efficiencies, which depend on the concentrations of functional proteins, remain approximately constant. This explains why linear modeling frameworks such as enzyme-constrained FBA and RBA, which assume constant enzyme efficiencies, provide a good approximation for idle protein expression.

Idle protein costs are additive: growth cost depends only on the combined mass fraction allocated to idle proteins, making different idle proteins interchangeable. This assumption is consistent with experiments showing additive growth costs for the expression of two fluorescent proteins (42).

### Trade-off between enzyme production and detoxification

While idle proteins impose a linear growth cost, functional proteins often impose indirect costs on a cell such as metabolic burden, membrane damage, protein misfolding, and other damaging effects (14, 24, 25). Such effects are particularly relevant during biotechnological production of heterologous compounds such as biofuels.

Turner *et al*. (24) showed that an intracellular accumulation of biofuel molecules can be toxic (i.e. decrease growth rate), but that this toxicity can be mitigated by overexpressing efflux pumps that export the compound. However, these pumps also cause a growth cost, and their optimal proteome fraction depends, potentially, on kinetic parameters of all the proteins in the cell. Overall, this means that cells must balance production of the toxic metabolite with the cost of detoxification.

To capture this trade-off, we extended our base model by adding two new reactions: one producing a toxic metabolite F from the carbon source, and another one exporting it from the cell:

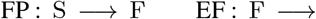

To model toxicity, we assume that the intracellular concentration of F inhibits protein translation by increasing the ribosome turnover time, with an inhibition constant 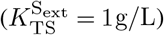.

To find the best cellular configuration, we varied the proteome fractions allocated to the biofuel-producing enzyme (*ϕ*_FP_) and the catalytic rate of the efflux pump 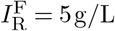, keeping 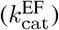 constant. The resulting optimum point depends on the 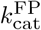 value, similarly to the experimental results in (24) (Figure 5). When the pump is assumed to be inefficient 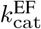, the model predicts that no efflux pumps should be produced – the cost of the pump exceeds its potential benefit (31).

**Fig. 5.**
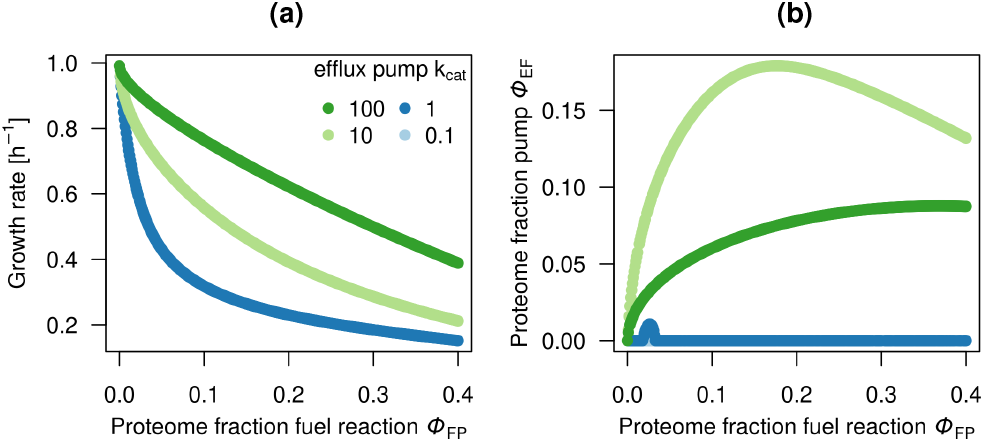
Trade-off between the production of an inhibitory product and the expression of an efflux pump. **(a)** A toxic product F inhibits ribosome activity, reducing growth. Growth can be improved by expressing an efflux pump, **(b)** with the optimal pump level depending on its turnover number 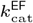.

This example shows that GBA can be used to study tradeoffs between different costly processes – here, toxic metabolite production and detoxification. Once parametrized with quantitative data, the model may guide the optimal expression of efflux pumps or other mitigating proteins. However, more detailed measurements would be required to make such predictions quantitative. The data from (24) report inducer concentrations but not the corresponding proteome allocations or intracellular biofuel levels, which hinders a direct parameterization of the model. Integrating proteomic and metabolomic data would enable more accurate studies of such trade-offs in future work.

### Transporter cost depends on the presence of alternative transporters

Finally, we used our model to study choices between alternative transporters for the carbon source. Experimental data from *Salmonella typhimurium* growing in maltose minimal medium (27, 43) show that reducing expression of the glucose transporter (PTS IIA^GLC^) to zero does not completely abolish growth; cells retain about 50% of their maximal growth rate. This indicates that alternative transporters can support maltose uptake, although with lower efficiency. We therefore used GBA to determine how the presence and efficiency of an alternative transporter affect the growth cost of the primary transporter.

We added a second transporter for the main carbon source (S_ext_), denoted TS2, into the base model (Figure 1). We then varied the proteome fraction of the main transporter TS1 while changing the kinetic parameters of TS2 (Figure 6). The concentration of S_ext_ was set to 10 g*/*L, ensuring that the primary transporter TS1 operates near saturation 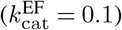.

**Fig. 6.**
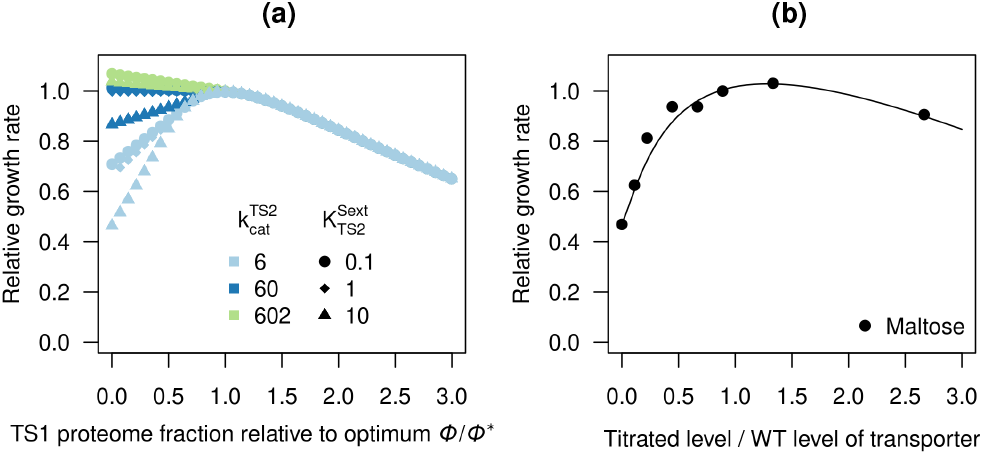
The availability of an alternative transporter reduces the costly effect of underexpression of the main transporter. **(a)** Relative growth rate as a function of the relative proteome fraction allocated to the primary transporter (TS1) for different catalytic efficiencies of the alternative transporter (TS2). When TS2 is efficient, it compensates for reduced TS1 expression; when less efficient, growth declines more sharply. **(b)**The points are experimental data from (43), redrawn from (27), showing the relative growth rate at different expression levels of transporter PTS IIA^GLC^ in *S. typhimurium* grown on maltose. The curve shows the phenomenological fit from (27).

The effects of TS1 underexpression depend on the kinetic properties of the alternative transporter TS2. When both transporters had identical *k*_cat_, reducing TS1 expression had no effect on growth, as the resulting upregulation of TS2 fully compensated for the reduction. When TS2 was less efficient, growth decreased progressively with lower TS1 expression. Notably, reducing the *k*_cat_ of TS2 by tenfold lowered growth to about 60% of the maximum when TS1 expression was zero, showing that these effects are nonlinear.

The growth cost of TS1 overexpression was largely independent of TS2 properties, because excess TS1 expression cannot be compensated by reducing the expression of the alternative transporter (which already has a zero expression level). Experimentally, a threefold overexpression of PTS IIA^GLC^ reduced the growth rate to approximately 85% of its maximum (27, 43), whereas the model predicted a reduction to approximately 65%. Because the growth cost is determined by the absolute increase in proteome allocation, the same relative level of overexpression can have different effects depending on the initial proteome fraction allocated to the transporter. However, a fully quantitative comparison is not possible because the absolute proteome allocation of PTS IIA^GLC^ was not measured experimentally.

This analysis demonstrates that GBA can capture the interplay between parallel reactions and quantify how kinetic differences shape their compensatory roles. Such models could, in principle, be used to estimate kinetic parameters of alternative or promiscuous enzymes from expression-growth data.

## Discussion

### Linking protein expression to growth with growth balance analysis

Our results demonstrate that GBA can be used to study the growth costs of protein over- and under-expression. By coupling enzyme kinetics, metabolite concentrations, and a proteome allocation constraint, GBA captures both the direct translational cost of protein synthesis and the indirect, non-linear system-level consequences that arise through flux redistribution and changes in intracellular metabolite concentrations. GBA belongs to a similar class of coarse-grained self-replicator models introduced by Molenaar et al. (1), which differ by being restricted to irreversible Michaelis-Menten kinetics and constrains only the total protein concentration. The GBA framework facilitates the creation and analysis of models in that class via an explicit function for the growth rate in terms of a minimal set of independent variables, which facilitates the derivation of the marginal value of each protein and its decomposition into four terms (31), the density constraint includes all biomass components so that metabolite accumulation reduces the space for proteins, and general rate laws such as reversible kinetics can be used (10).

GBA reproduces nonlinear relationships between suboptimal protein expression and cell growth in coarse-grained models, consistent with the experimental results for the suboptimal expression in actual proteins (11, 27) (compare Figure 2 with Figure S6). These protein-cost curves reflect specific kinetic parameters and pathway contexts in the present coarse-grained model, and hence may also differ across proteins in actual metabolic networks.

The concave dependence of growth rate on protein concentrations seen around the optimum in Fig. 2 agrees with a recent proof that growth rate is a concave function of enzyme concentrations in elementary flux modes (44). That proof assumes quasi steady state metabolism, a constant total protein concentration, and negligible metabolite mass and dilution by growth. GBA does not make these assumptions, so concavity does not hold for every protein in every condition. When metabolites accumulate, they take up part of the cellular density, and total protein decreases. The model therefore extends the concavity result to conditions where metabolite pools are not negligible.

For each reaction in the coarse-grained model, we obtain a corresponding optimal proteome allocation at which the growth rate is maximized, where the marginal benefits of increasing catalytic capacity exactly balance the marginal costs of additional proteome investment. Away from this optimum, costs outweigh benefits. Below the optimal fraction, insufficient enzyme abundance constrains the catalyzed flux. Above the optimum, allocating additional resources to the protein reduces growth, at least in part by displacing other cellular functions that are equally growth-limiting (10). This balance can be quantified by a breakdown of marginal costs and benefits (31) (Eq. (S1)), so the total marginal value of a protein emerges from different mechanisms, and depends on its context in the reaction network (Figure S1). These results emphasize that protein expression should be viewed in the context of the entire cellular network, where the effects of one enzyme propagate through metabolic interactions and competition for common resources.

This decomposition also reveals functional couplings between cellular subsystems. For example, increasing transcriptional capacity via RNAP raises the marginal benefit of nucleotide biosynthesis (NTS), because higher transcription flux increases demand for nucleotide supply. While such couplings are intuitive in small models, the same formalism applied to larger, more detailed networks could identify less obvious co-dependencies and guide rational co-expression strategies in metabolic engineering and synthetic biology.

Suboptimal protein expression can incur growth costs beyond those associated with transcription, translation, and molecular crowding. Several studies have shown that these costs vary across different classes of proteins, both in prokaryotes and eukaryotes (13, 14, 20, 21, 23, 40, 45). Multiple mechanisms may underlie these cost differences, some of which can be explicitly represented within the GBA framework.

One potential mechanism is metabolic burden: protein overexpression may cause an accumulation or depletion of intermediates, thereby disrupting fluxes. However, a systematic overexpression of glycolytic enzymes in yeast (one by one) led to only minor metabolomic changes, suggesting that metabolism can buffer such perturbations (14). Similarly, Tomala et al. (23) did not find a clear connection between enzyme function and its growth cost. Eguchi et al. (14) proposed that enzymes with reversible kinetics may buffer flux perturbations, allowing metabolism to re-equilibrate. When reversible kinetics are considered in our model, the predicted effects on growth rate and biomass composition decrease (compare Figure 2 and Figure S7). However, with irreversible kinetics, and in the lac model at high expression of the lactose enzyme, the model predicts that S or AA accumulate to a large fraction of the biomass, up to almost half (Fig. 3c and Fig. S9). Such accumulation is unlikely in real cells, which would limit it by regulation, for example by feedback inhibition or export, or growth would stop before this point. Adding an export reaction reduces the accumulation in the model (Fig. S11), so the growth costs predicted at high expression of such enzymes may be overestimated. In addition, some enzymes can be allosterically activated or inhibited – a phenomenon that can be included in GBA if a corresponding kinetic rate law is known.

In contrast to overexpressing native enzymes, overexpression of heterologous enzymes may cause more pronounced metabolic disruptions (46–48), as cells are not evolutionarily adapted to regulate these fluxes (47). An example is the mevalonate pathway for producing precursors for compounds including flavours, fragrances, pharmaceuticals, and biofuels, where the accumulation of HMG-CoA can inhibit fatty acid biosynthesis, leading to membrane stress and growth defects (49). Several terpenoids, including isoprenol, limonene, and linalool, have also been reported to inhibit growth in strains engineered to produce them. Proposed toxicity mechanisms include membrane disruption, interference with the proton motive force, and DNA damage (50).

GBA can in principle account for all these effects by including enzyme inhibition in the rate laws; in the simplest form, growth defects caused by a particular metabolite accumulation can be modeled by assuming that translation is inhibited by that metabolite. In more detailed models, specific mechanisms can be represented explicitly, for example by including the inhibition of ATP production through disruption of the proton motive force, or accounting for metabolite leakage.

Experimentally, toxicity can sometimes be mitigated by active export of toxic compounds. For instance, biofuels can be pumped out of the cell by efflux pumps, with these pumps introducing additional biosynthetic costs that depend on their kinetics (24). GBA can model the resulting tradeoffs (Figure 5), potentially guiding the selection and optimal expression of efflux pumps in bioengineering. We note, however, that our current implementation serves only as a simple demonstration: the toxic effect was modeled as a direct inhibition of translation. Quantitative predictions for real-world applications would require identifying the specific toxicity mechanisms and implementing appropriate kinetic rate laws with experimentally determined parameters.

Other types of burden, not yet captured by GBA, include protein misfolding, degradation, and regulatory feedback (14, 20, 23). Such effects can also make the cost of an idle protein nonlinear. Dekel and Alon (12) measured a nonlinear cost for the expression of the lac proteins in the absence of lactose, which may be related to the membrane transporter LacY, as discussed below. Further causes may include protein aggregation and signaling (20, 23). GBA instead predicts a linear cost, in agreement with other measurements (2, 19), because it describes an idle protein only by the space and resources that it uses. These effects could potentially be represented in future GBA models by incorporating rate laws that account for regulatory mechanisms – for example, metabolite-dependent activation of protein degradation, or inhibition of transport. However, experimental data are required to identify and quantify these effects, and to derive appropriate rate laws.

### Growth costs depend on environmental conditions

Environmental conditions influence whether the expression of specific proteins is costly or beneficial (12, 13). In line with this, our simulations approximating expression of the *lac* operon reproduce the general trends observed by Dekel and Alon: expression of a non-essential nutrient transporter reduces growth at low environmental substrate concentrations but can be beneficial at high substrate levels, when the use of this substrate as the main carbon source becomes economically advantageous for the cell (12).

However, the details of the cost function differ. Dekel and Alon reported a nonlinear, environment-independent cost, whereas our simulations predict environment-dependent costs. At low substrate concentrations in the environment, growth decreases roughly linearly with unnecessary protein expression, consistent with results by Scott et al. (2), who studied LacZ overexpression. At higher substrate concentrations, predicted growth costs of protein overexpression increase due to potential metabolic effects, such as the accumulation of intermediate metabolites. Whether such accumulation occurs *in vivo* remains unclear, as cells may buffer these effects through reversible reactions or enzyme inhibition.

The nonlinear costs observed by Dekel and Alon may also result from the expression of the entire *lac* operon, including the LacY transporter. Membrane proteins tend to cause higher growth costs than cytosolic proteins (20, 23), although LacY itself was not unusually burdensome in Stoebel et al. (51). In the presence of substrate, however, Eames et al. (52) showed that LacY-associated growth costs depend on both lactose and IPTG concentrations and are strongly influenced by LacY activity, likely through disruption of the proton motive force or substrate competition.

### Model limitations and future directions

We found that GBA predictions are close to experimental measurements, even with the simple models and rough parameter estimates used here. Achieving full quantitative accuracy would require more detailed experimental data. In particular, knowing the proteome fractions of expressed proteins, rather than inducer concentrations alone, would allow for a more precise model calibration (12, 24). To separate protein production costs from metabolic effects, GBA may be used to simulate mutations in the active sites of metabolic enzymes, as in (14), to quantify the cost of protein itself, distinct from metabolic or inhibitory effects.

GBA assumes an allocation of resources that is optimized for maximal growth, while empirical studies suggest that cells maintain a reserve capacity for protein synthesis (19). In baker’s yeast, for example, a protein can be overexpressed by up to 15% of the total proteome without affecting growth (14), implying that spare ribosomal capacity or downregulation of nonessential proteins can buffer the effects of over-expression. Similarly, *E. coli* produces many proteins that are not needed in the current environment (15, 16). Importantly, this unused protein capacity presents opportunities for biotechnology: a deletion of nonessential genes can increase growth rate (53) and redirect cellular resources toward a higher production of desired compounds (54–56).

GBA predicts the maximum growth rate by assuming an optimally allocated proteome in which all ribosome capacity is utilized, that is, without any spare capacity. However, the framework can be extended to account for spare capacity by introducing a lower bound on a nonfunctional protein mass fraction, thereby capturing the reduction in growth rate due to unused proteins. To evaluate whether maintaining spare capacity is advantageous under changing environments, dynamic simulations (57) would need to be performed, comparing two initial states: one with an optimally allocated proteome and one with reserve proteins. Upon environmental shifts (e.g., a switch in carbon source), cells with a fully optimized proteome may initially grow faster but acclimate more slowly, as new proteins must be synthesized. In contrast, cells with reserve capacity may respond more rapidly, having some proteins already available. Such simulations could quantify how temporary reductions in growth rate due to nonessential proteins are compensated by faster acclimation (15, 58, 59), potentially explaining why the cost of protein overexpression diminishes over several generations (16).

Dynamic GBA could also be extended to optimize the long-term average growth rate, rather than the instantaneous growth rate, under fluctuating environments. Such an approach could be used to investigate the evolutionary trade-off between maximizing growth in the current environment and maintaining reserve capacity or alternative metabolic strategies that improve long-term fitness (60).

Future GBA models could in principle provide more detailed growth cost estimations in specific contexts, for example by incorporating choices between alternative metabolic strategies (e.g., fermentation and respiration), several types of RNA, explicit degradation processes, and compartments such as mitochondria and nucleus for modeling eukaryotic cells.

Apart from overall proteome capacity, the limited surface area of the inner membrane imposes an additional constraint on membrane-embedded proteins such as transporters and ATP synthase. Membrane crowding has been proposed as a driver of overflow metabolism, as cells facing saturated membrane capacity reroute carbon through cytosolic pathways instead (61). In future work, this constraint could be incorporated into the GBA framework either phenomenologically, by imposing an upper bound on the membrane-protein proteome fraction (≈10% of proteome (61)), or more mechanistically by explicitly including lipid synthesis and enforcing a fixed lipid-to-protein area ratio within the membrane.

Genome-scale models that represent individual enzymes rather than aggregated pathways would enable a direct comparison with single-enzyme overexpression experiments. However, fully genome-scale models are currently intractable with this type of nonlinear optimization, although medium-scale models are feasible and extending the framework to larger models remains an important future direction. The main advantage of coarse-grained models like ours is their computational tractability, which enables systematic exploration of large parameter spaces and facilitates mechanistic interpretation while requiring less curation than larger-scale models (62). Despite their simplicity, such models reproduce a wide range of experimentally observed phenomena, including bacterial growth laws, realistic growth rates, and proteome reallocation during protein overexpression. Genome-scale or core metabolic models would enable more direct quantitative comparisons with experimental data, but at the cost of substantially greater computational complexity. In addition, the nonlinear optimization of GBA models is much harder than the linear optimization of enzyme-constrained models: the solution depends on the initial values, and the solver can fail for initial values far from the optimum (see Methods). Computational improvements, including better initialization strategies, convex approximations, and parallelization, will be important for extending nonlinear growth-cost analyses toward larger-scale models. A further limitation is that, unlike for enzyme-constrained models, for which tools such as GECKO (9) and AutoPACMEN (63) build models automatically from genome-scale reconstructions and kinetic databases, there is currently no automatic procedure to derive coarse-grained GBA models. They must be built manually, and their kinetic parameters must be estimated by an expert on a case by case basis.

Together, these results show that GBA provides a unified framework for linking protein expression to whole-cell physiology. Unlike approaches that describe protein costs through empirical functions, GBA derives these costs from resource-allocation and kinetic constraints and explains some important cell features from first principles such as (i) the biomass composition in optimal and suboptimal conditions, including metabolite concentrations; (ii) the intrinsic coupling of protein expression levels, where changes in one protein require compensatory changes in others; (iii) the dependence of protein costs at low or high expression on different cost factors (including metabolite burdens); (iv) the environment-dependence of how protein expression influences growth, and a clear indication that its effects cannot, in general, be separated into independent cost and benefit functions; and (v) the impact of toxic by-products and alternative substrate uptake reactions on growth and resource allocation. By connecting protein allocation, metabolism, and growth, GBA could also provide a mechanistic basis for guiding rational metabolic engineering strategies.

## Methods

### Growth balance analysis

Growth Balance Analysis (GBA) is a mathematical framework for modeling and analyzing nonlinear resource allocation models (10, 31). A GBA model is defined by a mass fraction matrix **M**, a multivariate function of reaction turnover times ***τ*** = ***τ*** (**a, c**) that depends on external and internal reactant concentrations **a, c**, and a cellular mass density *ρ*.

The total mass-fraction matrix **M**_total_ includes rows corresponding to external reactants and is directly related to the more conventional stoichiometric matrix **S**_total_ as follows. First, each row of **S**_total_ is multiplied by the molecular mass of the corresponding reactant. Then, each resulting column is normalized so that its positive entries sum to 1 and its negative entries sum to −1. The resulting matrix **M**_total_ therefore quantifies, for each reaction, the mass fraction of each reactant that is consumed (negative entries) or produced (positive entries). The matrix **M** is obtained from **M**_total_ by excluding the rows corresponding to external reactants.

For a given nonlinear cell model defined by (**M, *τ***, *ρ*) and given fixed external concentrations **a**, GBA focuses on the general problem of maximizing balanced cell growth rate *µ* (in units of 1*/*h) under the following constraints

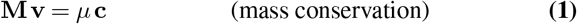

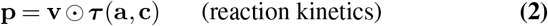

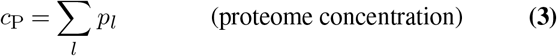

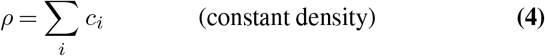

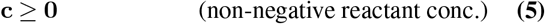

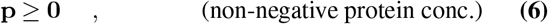

where **v** is a flux vector, **p** is a protein concentrations vector, and ⊙ indicates element-wise multiplication. Here the cell state depends on three different types of variables (**c, p, v**), and sensitivity or growth cost studies are not directly possible: changing one of these variables while keeping all other variables constant causes a violation of the constraints, since these are interdependent. In other words, there are no explicit partial derivatives of the growth rate with respect to any of these variables (**c, p, v**). We solve this problem in GBA by reformulating these constraints (Eqs. (1-6)) on a minimal set of dimensionless variables **q** (31), defined by normalizing the mass fluxes **v**, with units of g*/*(L h), with the growth rate *µ* and fixed density *ρ* with units of g*/*L

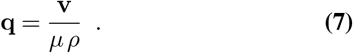

From the definition (Eq. (7)), we can rewrite the mass conservation constraint (Eq. (1)) as

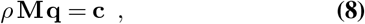

i.e., the internal reactant concentrations **c** satisfying mass conservation are automatically defined by **q**. This means the functions ***τ*** = ***τ*** (**a, c**) can now be understood as functions ***τ*** = ***τ*** (**a**, *ρ* **Mq**) of the variable **q** (and given constants **a**, *ρ*, **M**), and all the constraints (Eqs. (1-6)) can be compressed into a single objective function and constraints on **q** (31)

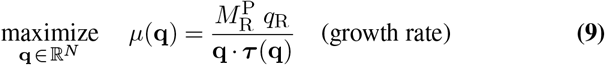

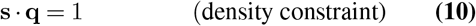

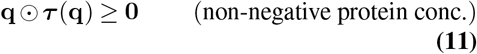

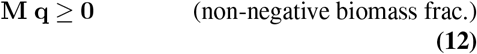

where 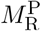 is the element in the last column (ribosome reaction) and last row (total protein) of the **M** matrix, *q*_R_ is the flux fraction of the ribosome reaction, the vector ***τ*** contains reaction turnover times, and the vector **s** denotes column sums of **M** (nonzero only for transport reactions).

The optimization yields the maximum growth rate *µ*^∗^ and the corresponding optimal flux-fraction vector **q**∗. Together, *µ*^∗^ and **q**∗ uniquely determine the complete balanced-growth state of the cell. The mass fluxes follow from Eq. (7), the concentrations of all metabolites and macromolecules from Eq. (8), and the protein concentrations from Eq. (14).

The problem can still be further simplified by incorporating the equality constraint (Eq. (10)) into the objective function (Eq. (9)), resulting in an optimization problem on truly independent variables, with a single objective function and inequality constraints (Eqs. (11,12)) only (31). This is however not necessary for the application in this study. Thus, effectively, we reformulated the problem on a much smaller set of independent variables **q**, and explicit partial derivatives of the growth rate *µ* with respect to each *q* can be calculated directly and represent closed expressions for the marginal value of each reaction, in terms of different types of marginal costs and benefits (31).

To model enzymes with reversible kinetics, we used convenience kinetics (64, 65) accounting for reversibility (i.e. product inhibiting the reaction rate), activation and inhibition with turnover time *τ*_*l*_ for the vector **x** = (**a, c**) of concatenated external and internal concentrations **a, c**, see Eq. (13).

Here, 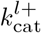 and 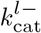 are the forward and backward catalytic rate constants of enzyme *l* (g*/*(g h)), 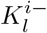 are the Michaelis constants for substrates (g*/*L) and 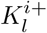 are the Michaelis constants for products (g*/*L). The constants 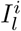 represent inhibition by reactant *i* in reaction *l*, while 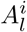 denotes activation by reactant *i* in reaction *l*.

This formulation provides a convenient framework that encompasses irreversible and reversible reactions and allows for the systematic inclusion of regulatory effects. In particular, irreversible kinetics are recovered by setting 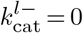 and taking 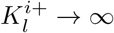. Reactions without activation or inhibition are obtained by setting 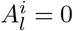 and taking 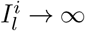. RNAP, DNAP, R were always assumed to be irreversible.

In the GBA formulation on variables **q**, protein concentrations **p** in units g*/*L are determined as (31)

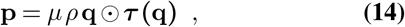

with corresponding proteome mass fraction of each reaction *l*

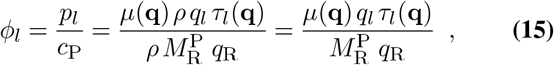

where *c*_P_ is the total proteome concentration also in g*/*L units, equal to 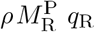 according to equation Eq. (8), and *µ*(**q**) is the function determined by equation Eq. (9). Thus, we may also understand now the proteome mass fractions ***ϕ*** = ***ϕ***(**q**) as functions uniquely determined by **q**.

### Modeling idle proteins

Idle proteins, including the non-growth-associated proteome (Q sector), are represented in the simulations shown in Figures 4 and S10 as a column of zeros in the **M** matrix. Consequently, the corresponding reaction neither consumes nor produces any metabolites. However, the protein still imposes a growth burden because it must be expressed when a lower bound is imposed using Eq. (16).

### Optimization

In this study we consider the balanced growth optimization defined by Eqs. (9-12), plus the inequality constraint

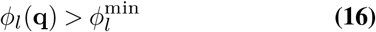

for the forced minimal over-expression 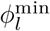 of a protein *l*, or the inequality

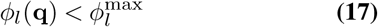

for the forced maximal under-expression 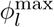 of a protein *l*.

The numerical solution of the corresponding nonlinear optimization problem was done using R version 4.6.1 (66), with the function `lp`from the package lpSolve (67), using the local solver SLSQP, local tolerance `localtol`set to 10^−13^ and relative tolerance `xtol_rel`set to 10^−10^. The numerical optimizations require initial starting values for the variable **q**, which were determined as described below.

### Estimating initial values for numerical optimization

For the general problem of finding initial values for the nonlinear optimization, we define the subproblem of finding feasible non-negative “flux fractions” **q** satisfying the linear optimization problem for a given vector **b**_min_ of minimal biomass fraction allocated to each reactant in the model

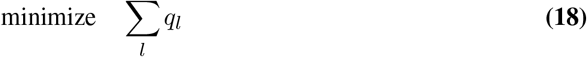

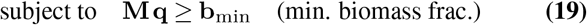

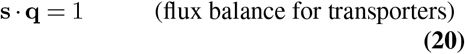

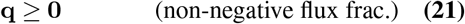

This linear optimization can be seen as a type of parsimonious FBA (68) on the variables **q** corresponding to irreversible flux fractions, with the lower bounds **b**_min_ emulating the usual biomass reaction. The flux balance constraint on transporters may also be seen as fixing the biomass production yield in the model. The linear problem was solved using the R function `lp` from the package lpSolve (67) and linear fits were performed with the function `lm`.

The solution of the linear optimization was used as the initial value for the nonlinear optimization in reference conditions, while for perturbations in this reference state (including by forcing under- or overexpression of a protein) we used by default the previous solution of the nonlinear problem as the initial condition in the next optimization. However, since the numerical solver could not always find a solution for the non-linear problem, we use different initialization strategies in the following priority order:

1. previously computed solution (if available),
2. estimated initial values A (see below)
3. estimated initial values B (see below).

If the computation fails to converge using option (1), we proceed to (2), and if necessary, to (3).

The two variants used to construct initial values were:

A. The lower bound on protein levels was iteratively increased in steps of 0.005, starting from half the experimental

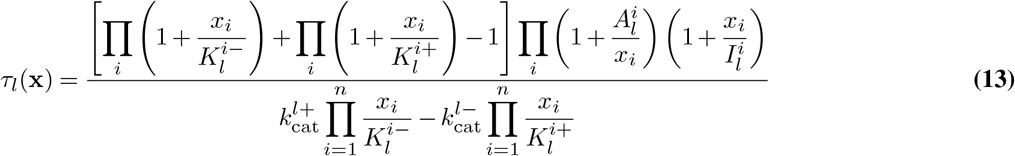

protein content (24.75%). For each tested protein content, if no feasible solution was found, the lower bounds **b**_min_ were reduced by a factor of 1.1 (up to 20 attempts) until a nonzero solution was obtained. Among all feasible solutions, the one yielding the highest growth rate according to Eq. (9) was selected.
B. The lower bound on protein level was fixed at the experimental value (55%). Lower bounds **b**_min_ were reduced by a factor of 1.1 (up to 20 attempts) until a nonzero solution was found.

### Parameters and experimental data

Kinetic parameters were estimated from fluxes and experimental proteomics data (see the next section, Eq. (23)).

Fluxes were estimated by linear optimization (see Section Estimating initial values for numerical optimization) where lower bounds on all biomass components are imposed through Eq. (19). These bounds are based on the experimental biomass composition of *E. coli* growing at of 1*/*h (BioNumbers ID 106154): 55% protein, 21% RNA, 3% DNA, 15% lipids and membrane polymers, 6% other. Since our coarse-grained model explicitly represents only the growth-associated proteome (assumed to be 45% of the total proteome (5)), RNA, DNA, and metabolites, these modeled components account for only about 55% of the cellular dry mass. Their biomass fractions were therefore rescaled to sum to 100%, giving 45% protein, 38% RNA, and 5% DNA. For metabolites, we assume biomass fractions of 1% for S, AA, NT and ATP, and 0.3% for ADP (the ATP/ADP ratio is roughly 3.2 in *E. coli*, BioNumbers ID 105021). These values were used as lower bounds in Eq. (19).

Protein abundance data were obtained from the *E. coli* proteomics datasets (69, 70). Protein abundances were mapped to model reactions using BioCyc annotations (71) (Table S2). Measurements from minimal-medium conditions in both studies were combined, fitted as a linear function of growth rate, and evaluated at *µ* = 1*/*h.

The biomass composition and proteomics data were used solely for parameter estimation. Once the kinetic parameters had been determined, all GBA simulations were performed without imposing biomass composition or protein abundance constraints. These quantities were instead predicted by the model, subject only to the constant biomass density constraint (Eq. (4)).

The dry biomass density of *E. coli* is approximately 340 g*/*L (BioNumbers ID 109049). Because the modeled biomass components account for only about 55% of the total dry mass, we scaled the density constraint to an effective value of 186 g*/*L in the simulations presented in the main text.

We also considered simulations including the non-growth-associated proteome (Q sector), assumed to constitute 55% of the total proteome (5) (Figures S10, 4). In this case, the modeled biomass components account for approximately 85% of the dry mass. After rescaling, the biomass fractions used for flux estimation were 65% protein, 25% RNA, and 4% DNA. The corresponding density constraint in the GBA simulations was scaled to 289 g*/*L.

### Estimating the kinetic parameters for the base model

Michaelis constants *K* were estimated based on the biomass concentrations **c** = *ρ* **Mq** determined by Eq. (8) with the initial values **q** estimated by strategy B (Section Estimating initial values for numerical optimization), which does not depend on kinetic parameters. For each reactant *i*, the Michaelis constant *K* was set to one third of its concentration *c*_*i*_ for all reactions using it as a substrate, and the *K*^+^ for products was set to be equal to the metabolite concentration. *K* for external metabolites S_ext_ and W_ext_ were set to 1 g*/*L.

Using flux fractions **q** predicted by the linear optimization (initial values B, Section Estimating initial values for numerical optimization), *E. coli* proteomics data at a growth rate of 1*/*h (69, 70) (see Table S2), we calculated the level of enzyme saturation in the general kinetic rate law (Eq. (13)) without activation nor inhibition as

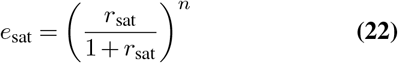

where *r*_sat_ is the ratio of metabolite concentration to *K*, set to value of 3, and *n* is the number of substrates. The transporter TS was assumed to be fully saturated.

The 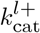 values (with reaction index *l*) were then estimated assuming the kinetics (Eq. (13)) without activation nor inhibition, so equation (Eq. (15)) results in

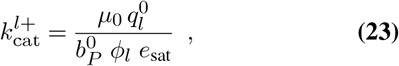

where 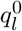 is the predicted flux fraction of reaction *l* in strategy B, 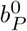 the corresponding protein fraction within the biomass, and *ϕ*_*l*_ is the experimentally measured proteome fraction mapped to reaction *l* at growth rate *µ*_0_ = 1*/*h (Table S2). Backward 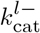 values were set to one fifth of the corresponding forward 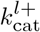.

With this initial estimation, GBA predicted a growth rate of 0.71*/*h. Subsequently, GBA was solved iteratively, adjusting all 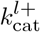 values by 5% upward or downward in each round until a growth rate of approximately 1*/*h was obtained.

Activation constants were set to the corresponding metabolite concentrations in the biomass. Specifically, 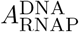 and 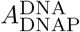 were set to 10 g*/*L, and 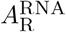 to 71 g*/*L.

In the model variant containing a LAC enzyme, the Michaelis constant of L_ext_ was set to 0.14 g*/*L (0.4 mmol*/*L, as in (12)). The turnover number of TS (glycerol utilization) was set to 20*/*h, and that of LAC to 15*/*h, as growth rate on lactose (0.53*/*h (72)) is roughly three-quarters of the growth rate on glycerol (0.69*/*h (35)). For LAC, we assumed irreversible kinetics. In simulations with an additional irreversible efflux of S, we set the *k*_cat_ value to an arbitrary value of 300*/*h and set *K* to 5.4 g*/*L (same as other reactions that use S).

We varied both the concentration of L_ext_ and the proteome fraction allocated to LAC (*ϕ*_LAC_), while keeping the primary carbon source S_ext_ (glycerol) at a subsaturating level of 0.57 *× K*. Note that the LAC reaction represents a lumped lactose transport and metabolism, and its parameters are not well-characterized. The concentration of glycerol relative to *K* was therefore fitted such that adding the second carbon source L_ext_ would provide the growth rate corresponding to the experimental observations at maximum L_ext_ concentration.

In simulations with toxic product production, the inhibition constant 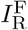 was set to 5 g*/*L and 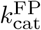 to 5*/*h, 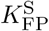 to 5.4 g*/*L (same as for other reactions using S), and 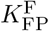 and 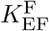 to an arbitrary value of 10 g*/*L.

## Data availability

All models (including parameters), code and data are available on GitHub (https://github.com/diana-sz/ProtCost) and archived on Zenodo https://doi.org/10.5281/zenodo.19683354 (73).

## ACKNOWLEDGEMENTS

This research was funded in whole or in part by the Austrian Science Fund (FWF) 10.55776/J4858.

## Supplementary Information

### Construction of the mass fraction matrix

GBA requires as input a mass fraction matrix **M**. It is constructed by writing down the reaction stoichiometry, multiplying each term by its molar mass (Table S1), and normalizing the result so that the mass fractions of the products sum to 1 and those of the substrates to -1.

Below we break down the construction of each reaction in a model. Note that the metabolite S represents a lumped pool of all cellular compounds that are not explicitly modeled.

#### ATPS

Cellular respiration can be represented with a reaction

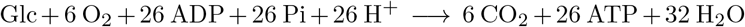

where we assumed 26 ATP molecules are made per glucose molecule (5). After lumping Glc, O_2_, Pi, H^+^ and H_2_O into S, and converting stoichiometric coefficients to mass fractions we obtain

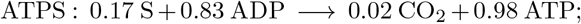

CO_2_ is labeled as W_ext_ in Figure 1.

#### AAS

We represented amino acid synthesis with a reaction

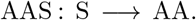

We assumed no ATP is needed for amino acid synthesis (74, 75).

#### NTS

The energy requirements for nucleotide synthesis were assumed to be 5 molecules of ATP per nucleotide (76). (The synthesis of dNTPs requires one more ATP, but since our model does not distinguish between NTPs and dNTPs, we used the value for RNA for simplicity.). The resulting reaction stoichiometry is

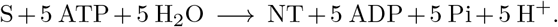

After lumping Pi, H^+^ and H_2_O into S and conversion to mass fractions we get

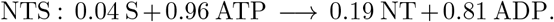

#### ADPS

For simplicity, we assumed that ADP can be synthesized from nucleotides

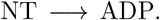

To account for the difference in masses, we added S as a product, yielding mass fractions

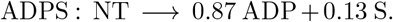

#### DNAP and RNAP

We assumed no ATP is needed for the polymerization of DNA and RNA as most energy is used for the synthesis of nucleotides (75, 77).

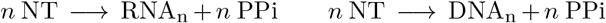

The diphosphate PPi released during polymerisation is represented as the general small compound S. After conversion to mass fractions, we get

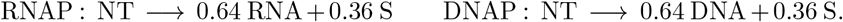

**R** Translation is represented with a reaction

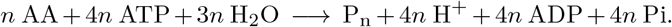

We assumed the polymerization cost of 4 ATP per amino acid (75, 76). Technically, making a protein of length n requires n-1 ATPs. However, assuming an average protein of 300 AA (BioNumbers 108984), we can neglect this for simplicity. After converting the stoichiometric coefficients into mass fractions and lumping the masses of H_2_O, H^+^ and Pi into S, we obtain

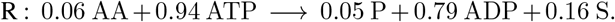

**Table S1.**
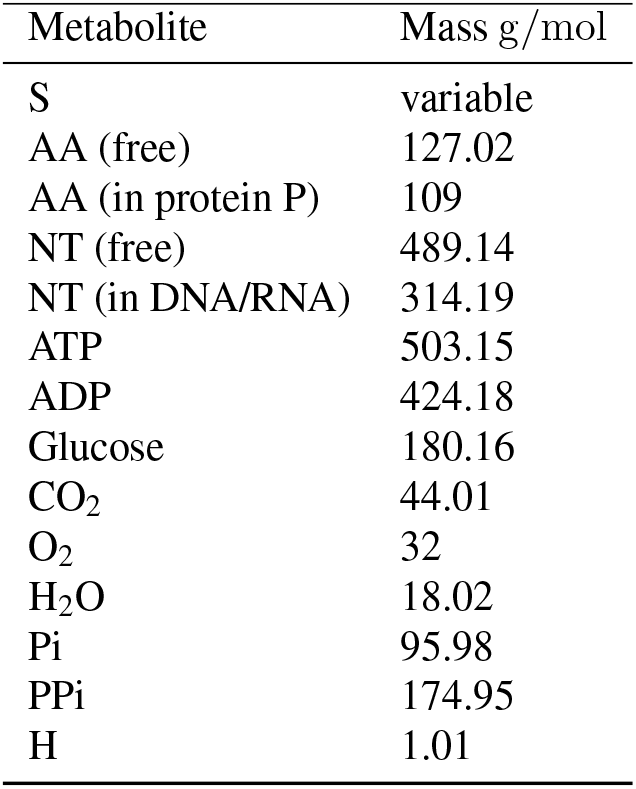
Molar masses used for constructing M.

| Metabolite | Mass g/mol |
| --- | --- |
| S | variable |
| AA (free) | 127.02 |
| AA (in protein P) | 109 |
| NT (free) | 489.14 |
| NT (in DNA/RNA) | 314.19 |
| ATP | 503.15 |
| ADP | 424.18 |
| Glucose | 180.16 |
| CO<sub>2</sub> | 44.01 |
| O<sub>2</sub> | 32 |
| H<sub>2</sub>O | 18.02 |
| Pi | 95.98 |
| PPi | 174.95 |
| H | 1.01 |

**Table S2.**
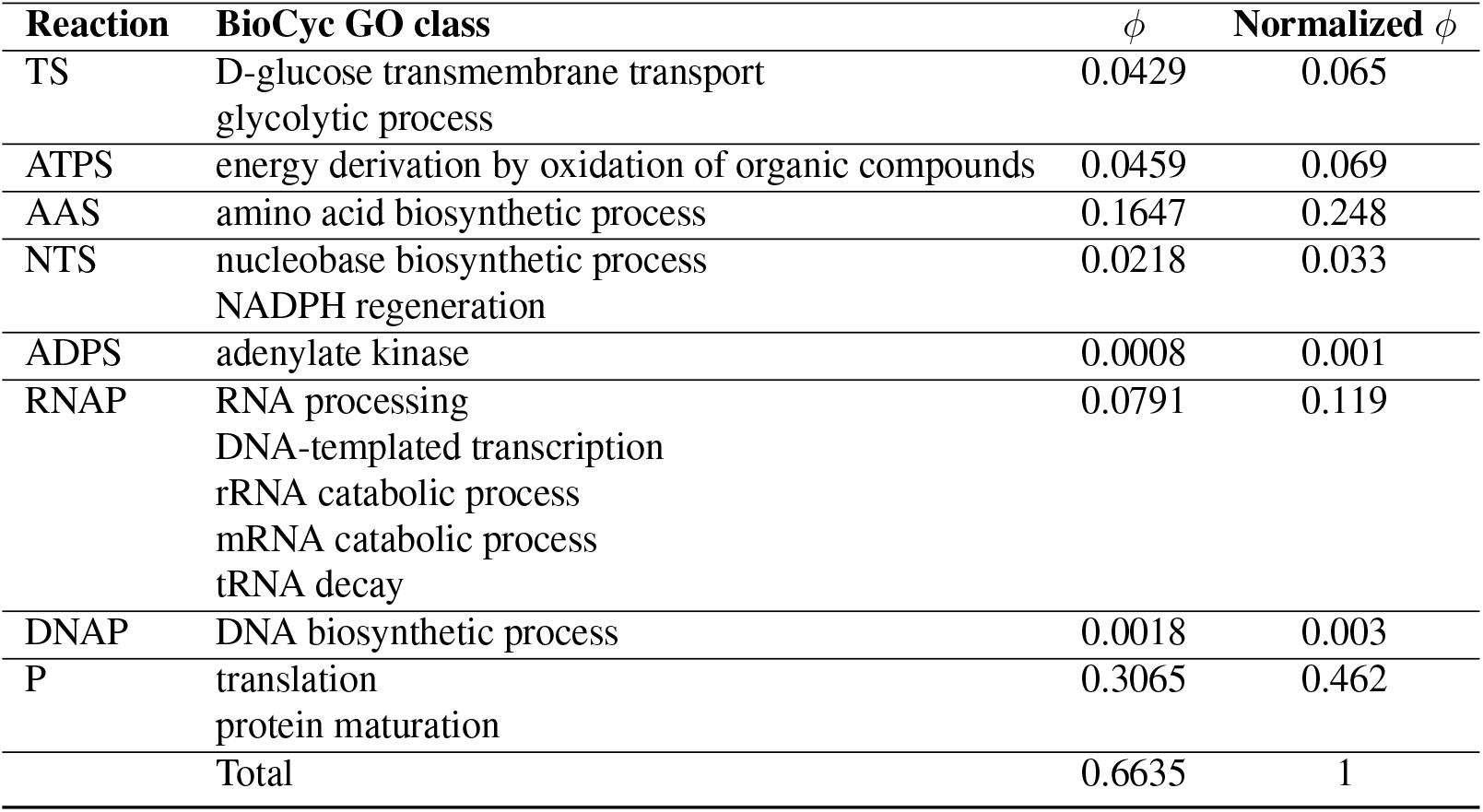
Mapping of *E. coli* proteomics data (69, 70) to the base model reactions (Figure 1). For each reaction, the corresponding BioCyc classes were identified, and protein mass fractions within each BioCyc GO class were summed. The resulting abundances were then normalized such that their total across all reactions equals 1. The normalized values were then used to estimate turnover numbers with Eq. (23).

| Reaction | BioCyc GO class | $\phi$ | Normalized $\phi$ |
| --- | --- | --- | --- |
| TS | D-glucose transmembrane transport<br>glycolytic process | 0.0429 | 0.065 |
| ATPS | energy derivation by oxidation of organic compounds | 0.0459 | 0.069 |
| AAS | amino acid biosynthetic process | 0.1647 | 0.248 |
| NTS | nucleobase biosynthetic process<br>NADPH regeneration | 0.0218 | 0.033 |
| ADPS | adenylate kinase | 0.0008 | 0.001 |
| RNAP | RNA processing<br>DNA-templated transcription<br>rRNA catabolic process<br>mRNA catabolic process<br>tRNA decay | 0.0791 | 0.119 |
| DNAP | DNA biosynthetic process | 0.0018 | 0.003 |
| P | translation<br>protein maturation | 0.3065 | 0.462 |
| Total |  | 0.6635 | 1 |

### Discussion on parameter estimation

The kinetic parameters were estimated based on experimental data from *E. coli* proteomics (Table S2) and biomass composition, and fluxes predicted from linear optimization (see Methods Section in the main text). This method of parameter prediction was chosen because the model has lumped reactions for which kinetic parameters are not readily available.

Another way to roughly estimate kinetic parameters would be to use the average reported *k*_cat_ for central carbon metabolism which is 78*/*s (78). This value needs to be converted to mass *k*_cat_ (mass of products per mass of enzyme per hour). Assuming an average enzyme molar mass of 40000 g*/*mol (BioNumbers ID 105861), and that roughly 100 enzymes participate in metabolism of each subsystem, we estimate mass-based turnover numbers of 9 g*/*(g h) for AAS, 937 g*/*(g h) for ATPS, and 217 g*/*(g h) for NTS. These estimates are of the same order of magnitude as values estimated in Table 2 in the main text. The larger values for NTS and ATPS compared to AAS arise because these reactions involve multiple ATP molecules, which contribute substantial mass.

### Changes in biomass composition upon enzyme overexpression

When assuming irreversible kinetics, overexpression of individual enzymes leads to substantial accumulation of certain metabolites (Figure S9). While this may seem straightforward, GBA predicts the allocation of the entire proteome, and in principle the cell could counteract such effects by adjusting the expression of other enzymes. We indeed find that biomass composition varies depending on which enzyme is overexpressed. For TS, AAS, ADPS and DNAP, overexpression results in pronounced accumulation of their respective products (S, AA, ADP, and DNA). In contrast, overexpression of NTS leads to the accumulation of RNA rather than nucleotides. This may be because RNA is required for ribosome function, so its accumulation is less detrimental to growth than the accumulation of free nucleotides. In the model with reversible kinetics (Figure S8), both nucleotide and RNA fractions remain approximately constant upon NTS overexpression. Interestingly, when RNAP is overexpressed, RNA levels remain roughly constant above *ϕ* = 0.2, likely because RNA synthesis depends on DNA, and the effect of increased RNAP is counterbalanced by downregulation of DNAP.

Overall, these results show that the effects of protein overexpression on biomass composition are nontrivial and depend on the kinetics and regulation of all other enzymes in the cell.

### Discussion on the definition of cellular density

In all simulations, we impose an upper bound on the total biomass density expressed as mass per unit volume (g/L), such that each cellular component contributes according to its mass. This approximation ignores that molecules with the same mass can occupy different physical volumes, so mass density is only an indirect proxy for intracellular crowding. In principle, one could instead constrain the total occupied volume of cellular components but this would require reliable estimates of molecular volumes, which are not always available. Since mass and volume are approximately proportional (79), we use a mass-density constraint as a practical approximation. Nevertheless, the density constraint could be easily adapted to account for a limited volume instead of mass.

### The protein cost and benefit contributions of a reaction

To understand how protein costs and benefits arise, we apply an analytical expression, derived previously in the context of GBA (31) for the total marginal value *V*_*l*_ of each reaction *l*. The marginal value is defined as the proportional impact a small increase in *q*_*l*_ has on the growth rate (scaled by the proteome mass fraction in biomass *b*_P_). In GBA models, it is given by

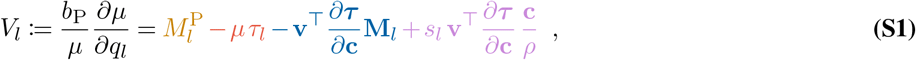

where *µ* is the growth rate (1*/*h), 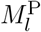 is the last row of the **M** matrix (quantifying how much of each flux is going into proteome synthesis), *τ*_*l*_ is the turnover time of reaction *l* (hour) determined by its kinetics (see Methods), **v** is the vector of fluxes, *s*_*l*_ is the sum of column *l* in the **M** matrix (nonzero only for transporters), **c** is the vector of reactant concentrations (g*/*L), and *ρ* is the cell density (g*/*L).

The relation (Eq. (S1)) shows how the total “economic value” *V*_*l*_ of each reaction *l* results from four different types of marginal costs and benefits (different colors), originating from separate impacts a small increase in *q*_*l*_ has on the growth rate via changes in proteome allocation. These separate cost and benefit contributions can be interpreted as follows (see mathematical details in (31)):

- Protein synthesis benefit 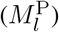 – the contribution of reaction *l* to increasing the growth rate via an increase in protein synthesis, which in turn provides more proteins available to catalyze reactions. In this framework, the ribosome reaction is the only one that synthesizes proteins, so it is also the only reaction with a non-zero synthesis benefit. This value quantifies effects mediated by the following causal chain: higher *q*_R_ *→* higher *c*_P_ *→* higher *µ*.
- Protein catalytic/catalyst cost (−*µτ*_*l*_) – the growth penalty associated with the diverted proteome necessary to catalyze the increased flux *q*_*l*_, thus reducing the proteome available to other reactions. This value quantifies the effects mediated by the following causal chain: higher *q*_*l*_ *→* higher *p*_*l*_ *→* lower *µ*.
- Protein saturation value 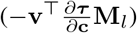 – the local impact of a reaction on increasing (positive value, corresponding to a benefit) or decreasing (negative value, corresponding to a cost) the necessary protein allocation to its surrounding reactions (including itself), via changes in their saturation due to changes in the concentration of their shared reactants, itself forced by mass conservation when increasing *q*_*l*_. This value quantifies the effects mediated by the following (more indirect) causal chain: higher *q*_*l*_ *→* changes **c** *→* changes ***τ*** *→* changes **p** *→* changes *µ*.
- Proteome crowding value 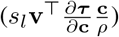 – the marginal impact of a transport reaction (the only reactions with *s*_*l*_ ≠ 0) on the biomass fraction available for proteins; a negative proteome crowding value indicates an import of mass, which decreases the available biomass fraction *b*_P_ for proteins and decreases growth rate, while a positive value indicates an export of mass causing the opposite effect. This value quantifies the effects mediated by the following causal chain: higher *q*_*l*_ *→* changes *ρ →* changes *c*_P_ *→* changes *µ*.

At optimal steady growth states, small changes on each *q*_*l*_ do not impact the growth rate and the marginal values *V*_*l*_ are zero, which means these four terms must balance exactly and sum to zero for every reaction. From an economic point of view, at the optimum the marginal costs of proteome allocation must be precisely compensated by their marginal benefits.

Table S3 shows this balance, for each of the catalytic proteins, in the state of maximal growth for the base model at a saturating environment, where all enzymes are expressed at their optimal levels. For most metabolic and information-processing enzymes (AAS, NTS, ADPS, RNAP, DNAP), the negative catalytic costs are exactly offset by equal positive local benefits (saturation value). The magnitudes of these terms reflect how strongly each reaction influences growth rate, with AAS having the largest impact and ATPS the smallest.

The crowding value is nonzero only for the transport reactions ATPS and TS and captures their indirect effects on biomass production. TS has a negative crowding value because higher intracellular concentration of S occupies more space in the cell, but this is compensated by its positive saturation value. In contrast, ATPS shows a positive crowding value because the secretion of waste frees up space in the cell.

The ribosome is unique in providing a synthesis benefit, balanced by its catalytic cost and negative saturation value, consistent with its high proteome investment.

Hence, proteins contribute to growth through three separate types of effects – directly by producing biomass, locally by enabling efficient flux, or indirectly through metabolite-mediated effects – and for optimal proteome allocation at steady state, these contributions must balance out. In turn, deviations from this balance (e.g., when expression deviates from the optimum) indicate that a protein is under- or overexpressed relative to the growth-optimal state. A positive marginal value implies that increasing the protein’s allocation would improve growth, while a negative marginal value implies that reducing its expression would be beneficial.

**Table S3.** The different protein costs and benefits of reactions in the base model at the optimal growth on saturating external substrate 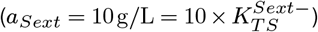. The four terms of Eq. (S1) balance to zero for each protein, as required at optimal balanced growth.

| Reaction | Synthesis benefit | Catalytic cost | Saturation value | Crowding value | Marginal value |
| --- | --- | --- | --- | --- | --- |
| TS | 0 | -0.023 | 0.231 | -0.207 | 0 |
| ATPS | 0 | -0.001 | -0.003 | 0.004 | 0 |
| AAS | 0 | -0.166 | 0.166 | 0 | 0 |
| NTS | 0 | -0.002 | 0.002 | 0 | 0 |
| ADPS | 0 | -0.007 | 0.007 | 0 | 0 |
| RNAP | 0 | -0.111 | 0.111 | 0 | 0 |
| DNAP | 0 | -0.020 | 0.020 | 0 | 0 |
| R | 0.05 | -0.026 | -0.024 | 0 | 0 |

### Decomposition of the (marginal) costs and benefits of reactions under protein under- and overexpression

We used Eq. (S1) to understand how protein under- and overexpression affects cell growth through four contributions: synthesis benefit, catalytic cost, saturation value, crowding value (Figure S1). For clarity, synthesis benefit is not shown because in our model, it is positive for the ribosome reaction R and zero for all other reactions.

For each protein, the marginal value (black curve) crosses zero at the growth-maximizing expression level (gray dashed line). Its behavior is straightforward to interpret along the diagonal. For example, when AAS is underexpressed, the marginal value is positive, indicating that higher expression would increase growth rate; when overexpressed, the marginal value becomes negative, indicating that reduced expression would be beneficial. Decomposing the marginal value reveals the causal chain underlying growth reduction. Upon protein overexpression, both the catalytic cost and the saturation value decrease, but the saturation value declines much more rapidly. This indicates that the growth reduction is driven primarily by global changes in intracellular concentrations, rather than by the energetic cost of synthesizing the excess protein alone. When a protein is underexpressed, its marginal value increases mainly due to an increase in saturation value, which can be interpreted as missing catalytic benefits resulting from insufficient enzyme abundance.

Next, we examine how the costs and benefits of other proteins respond to these perturbations. For example, if RNAP is overexpressed, the marginal value is positive for NTS and DNAP, suggesting the cell may benefit from co-expression of these proteins. Overexpression of NTS would benefit from co-expression of TS and ATPS. Similar behavior is seen for other proteins: local costs and benefits depend on which protein is perturbed.

These results show that the marginal costs and benefits of the different proteins, after a forced change of one of the protein levels, depend on which protein is under- or overexpressed. This means that these costs cannot be interpreted in isolation but must be analyzed in detail and in the context of the entire cellular network. This analysis may reveal functional dependencies between proteins by identifying what other proteins should be up- or downregulated along with the target protein. Such information can guide targeted co-expression strategies, for example in strain design or synthetic perturbations, where coordinated expression is required to improve efficiency.

We also tested how the different cost and benefit (Eq. (S1)) change with the environment in the LAC overexpression simulations (Figure S11 (c)). The catalytic cost is highest at low external lactose concentrations, where the LAC proteins are inefficient. At higher concentrations, the proteins become more saturated and therefore more efficient: *τ* ^LAC^ decreases, and therefore the cost decreases too. This holds for any tested value *ϕ*_LAC_. The saturation value depends strongly on *ϕ*_LAC_ and L_ext_ and largely determines the total marginal value. In both cases, it increases with L_ext_ concentration. For *ϕ*_LAC_ = 0.04, the saturation value is close to zero at low lactose concentrations, so the marginal value is dominated by the catalysis cost. As L_ext_ increases, the saturation value becomes positive and eventually outweighs the catalytic cost, leading to a positive marginal value. For *ϕ*_LAC_ = 0.3, the saturation value also increases with L_ext_ but it remains negative for all lactose concentrations. The benefits never outweigh the costs, so the marginal value is always negative, meaning expression is excessive regardless of L_ext_ concentration. Altogether, these results show that in some conditions the growth rate is primarily limited by the direct catalytic cost of protein expression, whereas in others it is governed by global cellular rearrangements captured by the saturation value.

**Fig. S1.**
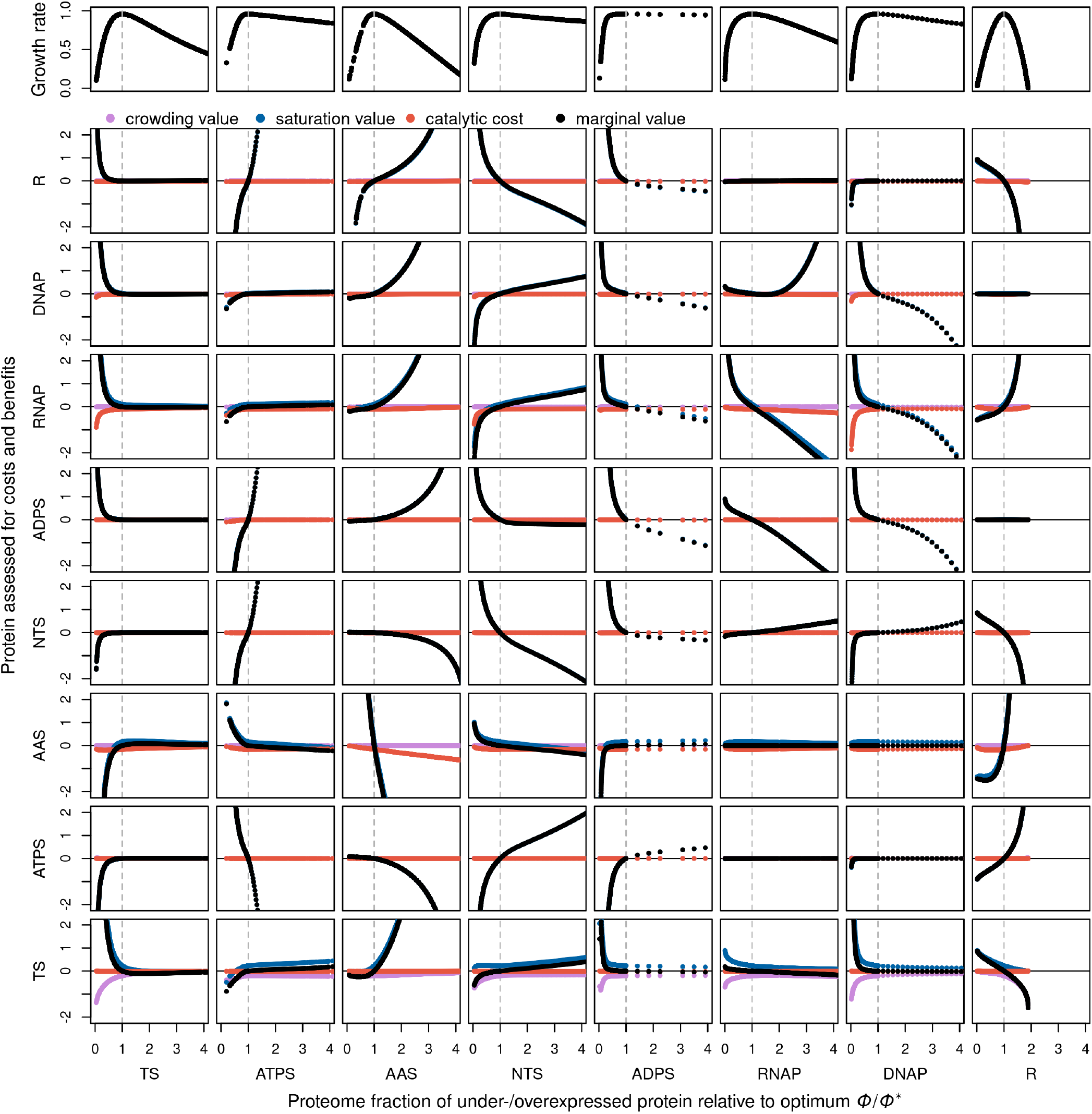
Decomposition of costs and benefits during protein perturbations, assuming reversible kinetics. Each column represents one protein perturbation (corresponding to the rows in Figure 2). The top row shows the growth rate, and the remaining rows show the decomposed cost and benefit terms for each protein (Eq. (S1)). The black curve is the total marginal value (the sum of the four terms), which crosses zero at the growth-optimal expression level. For each perturbed protein, its own marginal value is a decreasing function.

### Protein cost in a toy model

We tested whether GBA can capture the nonlinear effect of protein cost on growth rate using a minimal toy model, comprising only three reactions: uptake of the external nutrient S_ext_ by the transporter TS to produce the internal metabolite S, conversion of S into amino acids (AA) by the amino acid synthesis enzyme (AAS), and protein (P) synthesis by the ribosome (R). P represents the total modeled proteome, defined as the sum of all proteins in the model. The concentration of the external nutrient S_ext_ was set to 100 g*/*L. Despite its simplicity, the model reproduces the general trends observed in the larger model (Figure S2).

**Fig. S2.**
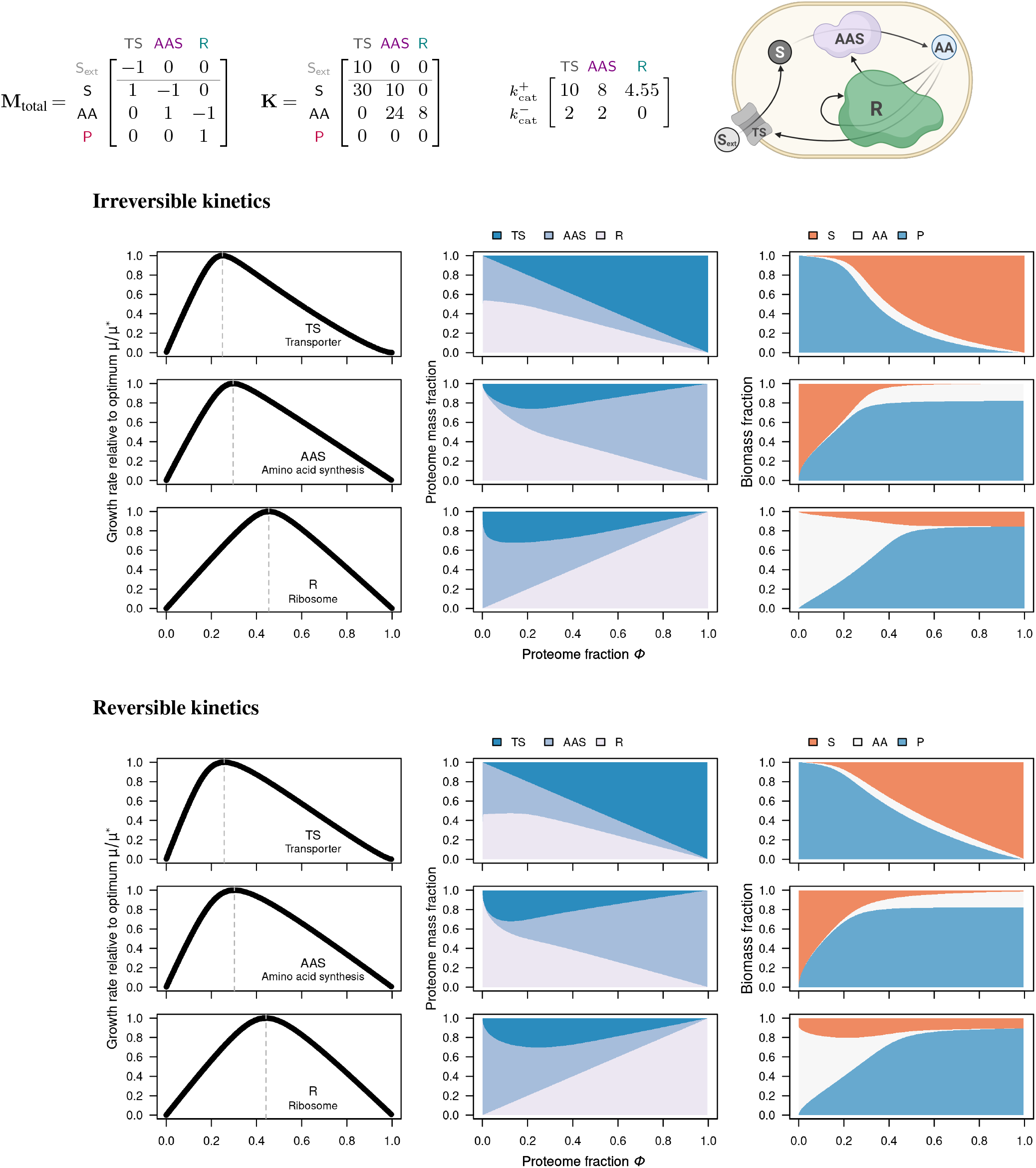
Growth costs in a toy model with three reactions with irreversible and reversible kinetics. The first column shows the normalized growth rate as a function of absolute *ϕ*. The second column shows the proteome composition, and the third the biomass composition. Cell figure created with biorender.com.

### Comparison with other modeling approaches

We compared the growth cost of under- and overexpression in GBA with three related approaches on the toy model with three reactions (TS, AAS, R), using the same turnover numbers and Michaelis constants (if applicable).

i. Enzyme-constrained FBA was computed with the COBRA-k package (39): each flux is limited by the product of the turnover number and the enzyme concentration, the sum of the enzyme masses is limited by the protein fraction of the biomass, and the biomass composition is fixed by a biomass reaction.
ii. The kinetic COBRA-k model adds the saturation term of the rate laws (reversible), with the metabolite concentrations as free variables constrained by lower and upper bounds (lower bound: 1 *×* 10^−6^ mol*/*L, the default of the package; upper bound: concentration of the species in the biomass; the concentration of the external carbon source is fixed). Thermodynamic terms were not included, because standard Gibbs energies are not defined for the toy model.
iii. RBA was formulated as in (38), with a capacity constraint for each enzyme, a ribosome capacity constraint that includes the synthesis of the ribosome itself, and a dry mass constraint. The maximal growth rate was found by bisection, using a linear program that tests at each growth rate whether a feasible solution exists.

The density (340 g*/*L), the external concentration of the carbon source (100 g*/*L), and the biomass composition (S 0.150, AA 0.0725, protein 0.777 g*/*gDW) were taken from the growth optimum of the GBA model. As in the main text, the proteome fraction of one enzyme was fixed to values between 0 and 1, and the growth rate was maximized over all other variables. A Jupyter notebook with all three methods is available on GitHub (https://github.com/diana-sz/ProtCost) and archived on Zenodo (73).

RBA and enzyme-constrained FBA give identical curves (up to numerical tolerances), because both have linear capacity constraints and a fixed biomass composition (Fig. S3). The curves consist of two straight segments with a sharp kink at the optimum. The kinetic COBRA-k model has a rounded optimum, but away from it, its curve is closer to the linear methods than to GBA. In COBRA-k, the biomass composition is fixed, and the metabolite concentrations are free variables within bounds. They affect the fluxes through the saturation terms, but, unlike in GBA, the fluxes do not determine the concentrations through mass conservation and dilution by growth. There is also no density constraint that couples proteins, metabolites, and macromolecules, because the enzyme pool is a separate bound. COBRA-k has an optional limit on the summed concentration of small metabolites, which we did not use. As a result, the effect of the enzyme levels on the saturation, and thus the rounded shape, is only partly captured.

**Fig. S3.**
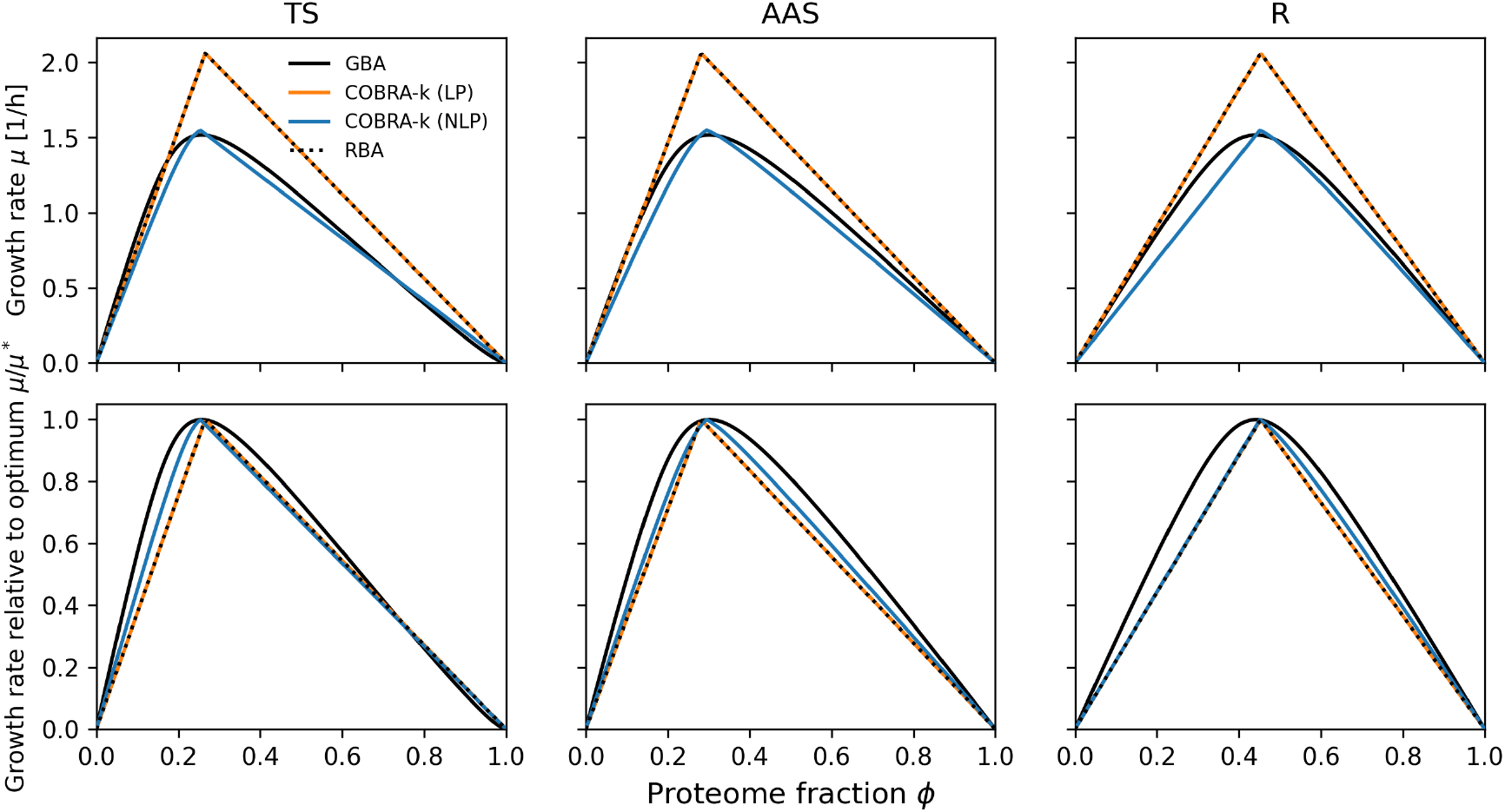
Growth cost of suboptimal expression in the toy model with GBA, enzyme-constrained FBA (COBRA-k LP), the kinetic COBRA-k model (COBRA-k NLP), and RBA. GBA and COBRA-k NLP are shown with reversible kinetics. Top: growth rate as a function of the proteome fraction *ϕ* of each enzyme. Bottom: growth rate relative to the optimum of each method. The proteome fraction of one enzyme is fixed, and the growth rate is maximized over all other variables. The curves of RBA (dotted) and of enzyme-constrained FBA (COBRA-k LP) coincide.

## Supplementary Figures

Additional figures related to the main text.

**Fig. S4.**
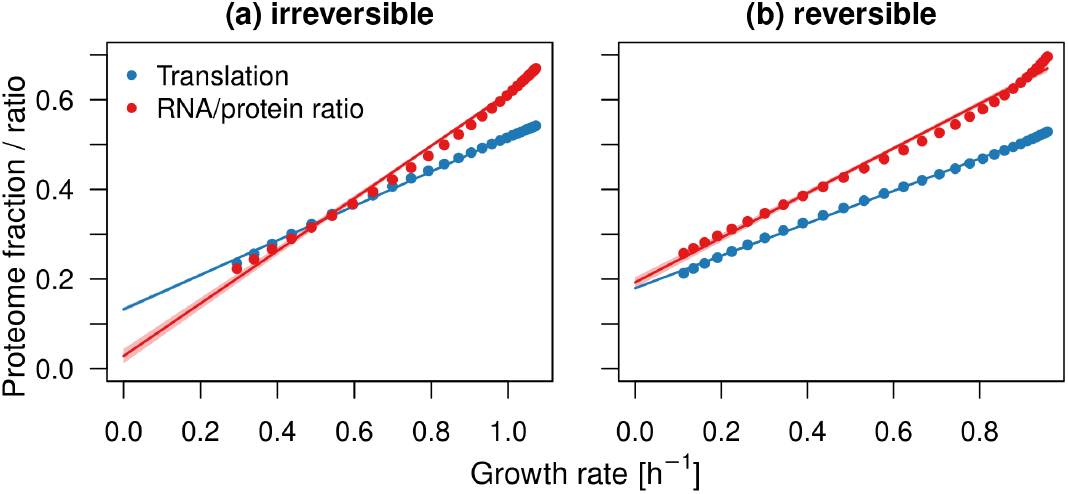
Validation of the model. The base model (Figure 1) reproduces the linear relationship of growth rate with the fraction of proteome associated with translation and with RNA/protein ratio (2). Growth rate was varied by changing the external nutrient (S_ext_) concentration.

**Fig. S5.**
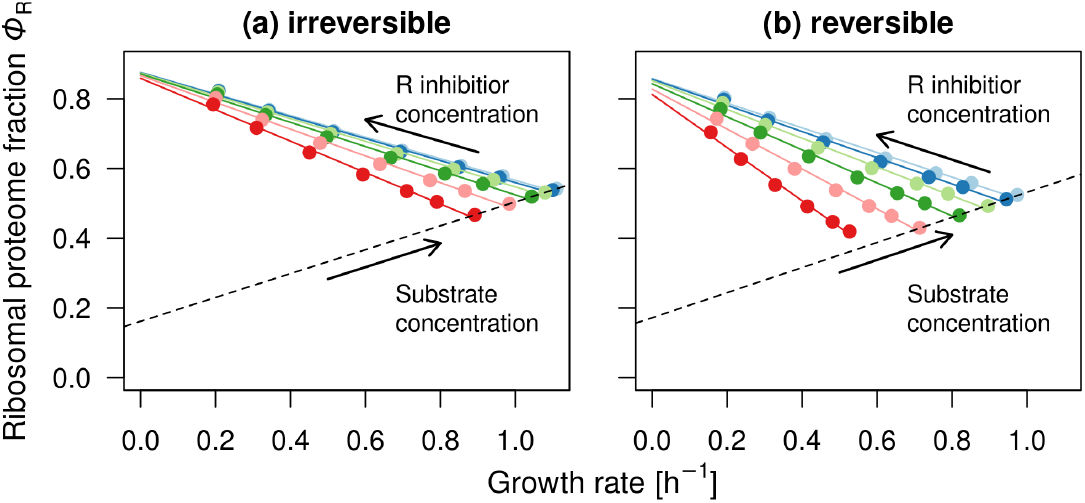
Base model captures the reverse growth law – increasing concentrations of a ribosome inhibitor reduce the growth rate while increasing the ribosomal proteome fraction. The ribosome inhibitor is modeled as an extracellular compound that acts directly on the ribosome and is incorporated into the ribosomal rate law as an inhibitor, with 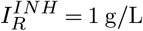 and inhibitor concentrations ranging from 0 to 10 g*/*L.

**Fig. S6.**
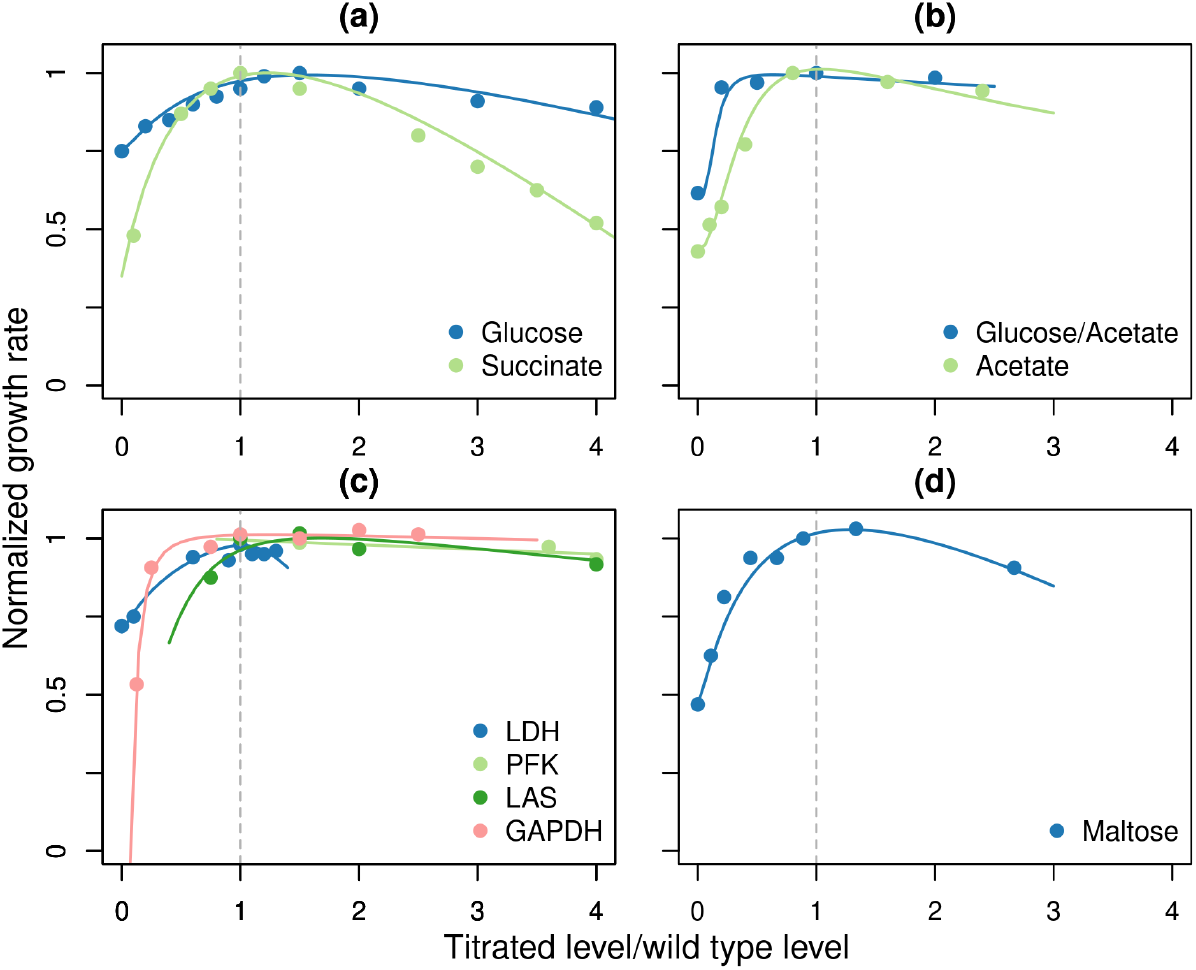
Experimentally measured growth cost of protein under- and overexpression. (A) ATP synthase in *E. coli* (80, 81); (B) citrate synthase in *E. coli* (82); (C) glycolytic enzymes in *Lactococcus lactis* (83–85); (D) PTS transporter system (glucose-specific subunit IIA) in *Salmonella typhimurium* (43). All proteins were expressed under control of an IPTG-inducible promoter. Figure redrawn from (27, 86).

**Fig. S7.**
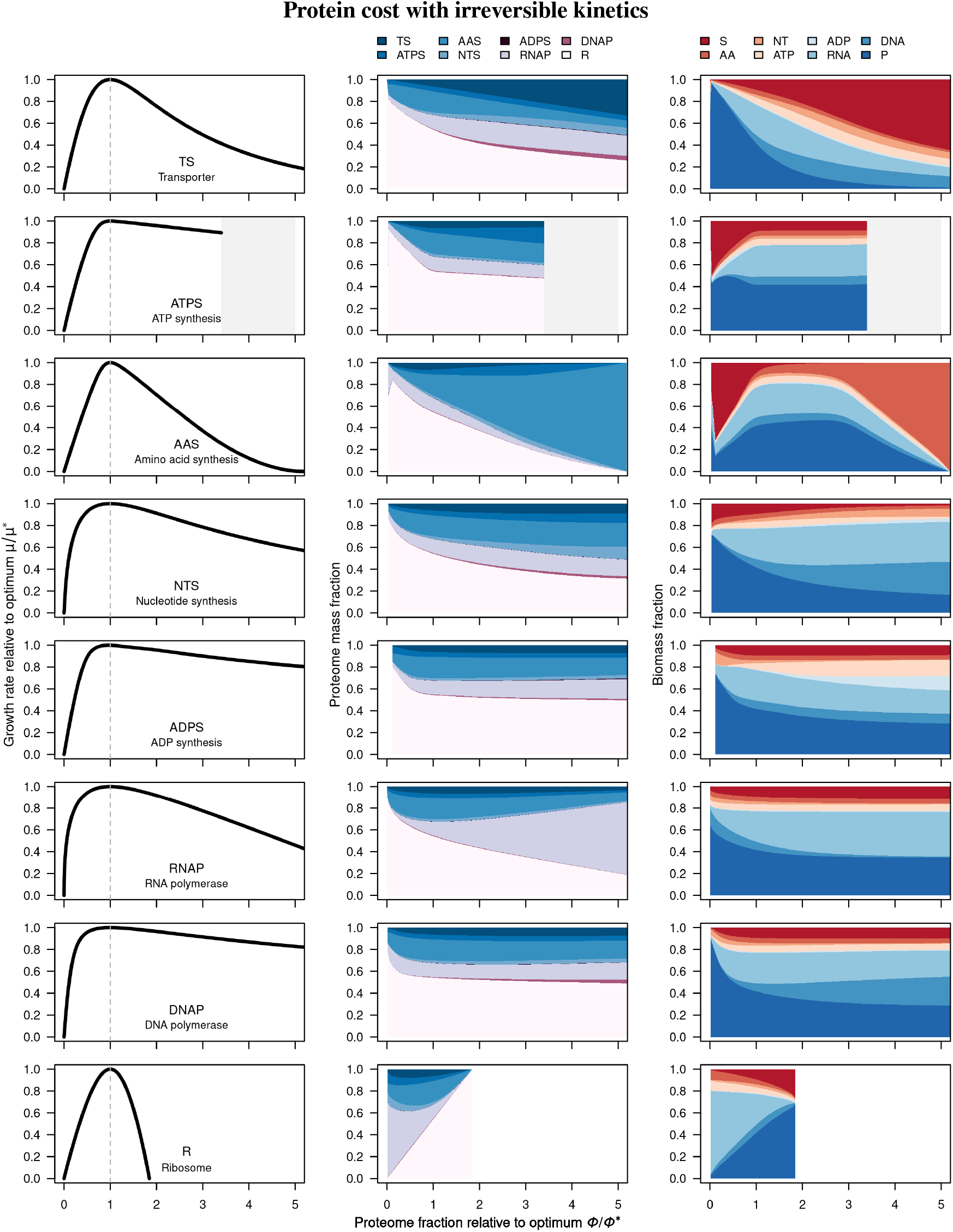
Effects of over- and underexpression of different proteins on growth rate, proteome composition, and biomass composition with irreversible kinetics. With irreversible kinetics, biomass composition is affected more than with reversible kinetics. The first column shows the normalized growth rate as a function of *ϕ* relative to the optimal *ϕ*∗. The second column shows the proteome composition, and the third the biomass composition. Numerical results for ATPS were not found for higher proteome fractions (indicated with the light gray region).

**Fig. S8.**
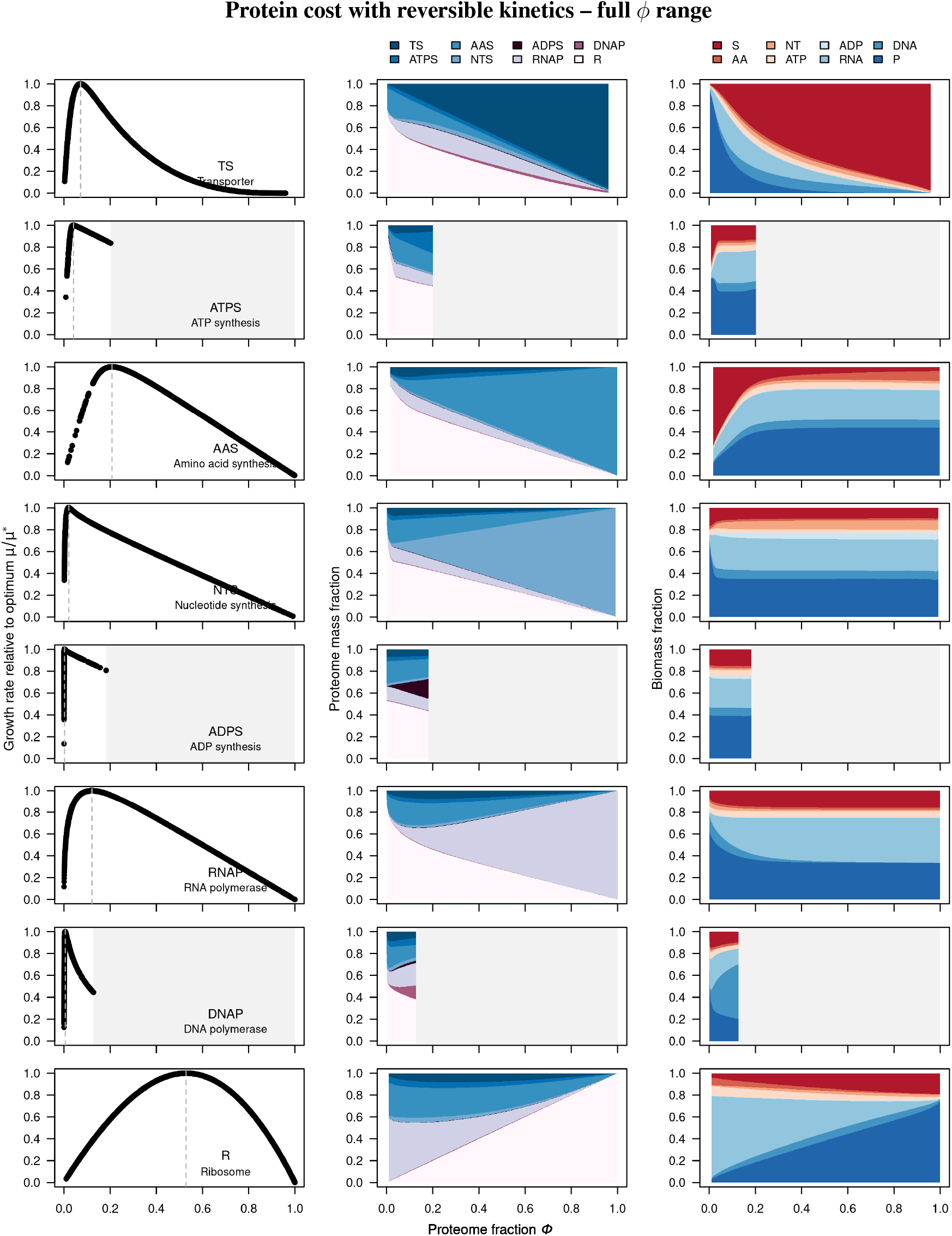
Effects of over- and underexpression of different proteins on growth rate, proteome composition, and biomass composition (reversible kinetics). The first column shows the normalized growth rate as a function of absolute *ϕ*. The second column shows the proteome composition, and the third the biomass composition. Note that for some proteins, simulations do not converge if proteins are overexpressed too much above or below the optimum. The figure corresponds to Figure 2 in the main text but shown over the full *ϕ* range. Numerical results for ATPS, ADPS and DNAP were not found for higher proteome fractions (indicated with the light gray regions).

**Fig. S9.**
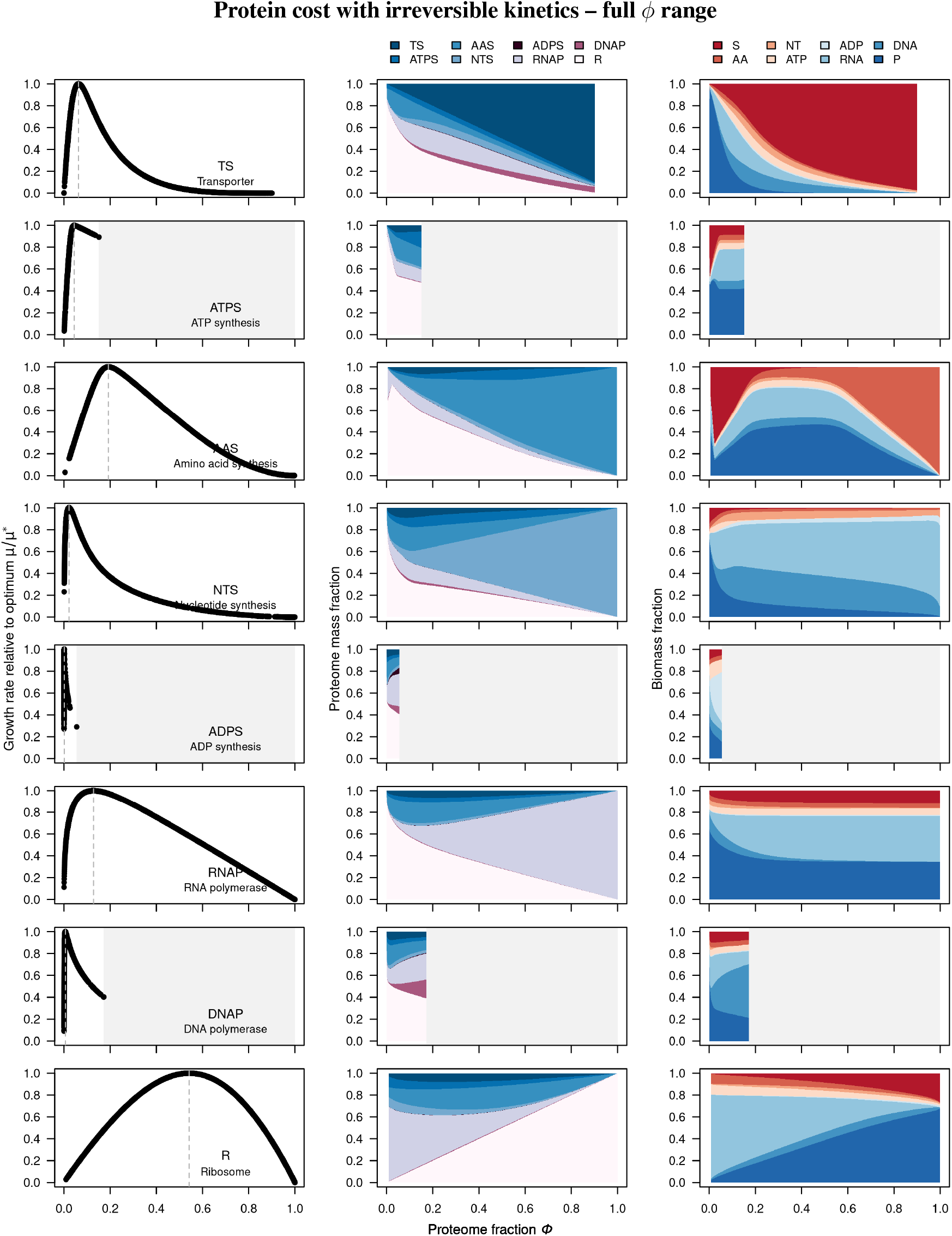
Effects of over- and underexpression of different proteins on growth rate, proteome composition, and biomass composition (irreversible kinetics). The first column shows the normalized growth rate as a function of absolute *ϕ*. The second column shows the proteome composition, and the third the biomass composition. Note that for some proteins, simulations do not converge if proteins are overexpressed too much above or below the optimum. The figure corresponds to Figure S7 but shown over the full *ϕ* range. Numerical results for ATPS, ADPS and DNAP were not found for higher proteome fractions (indicated with the light gray regions).

**Fig. S10.**
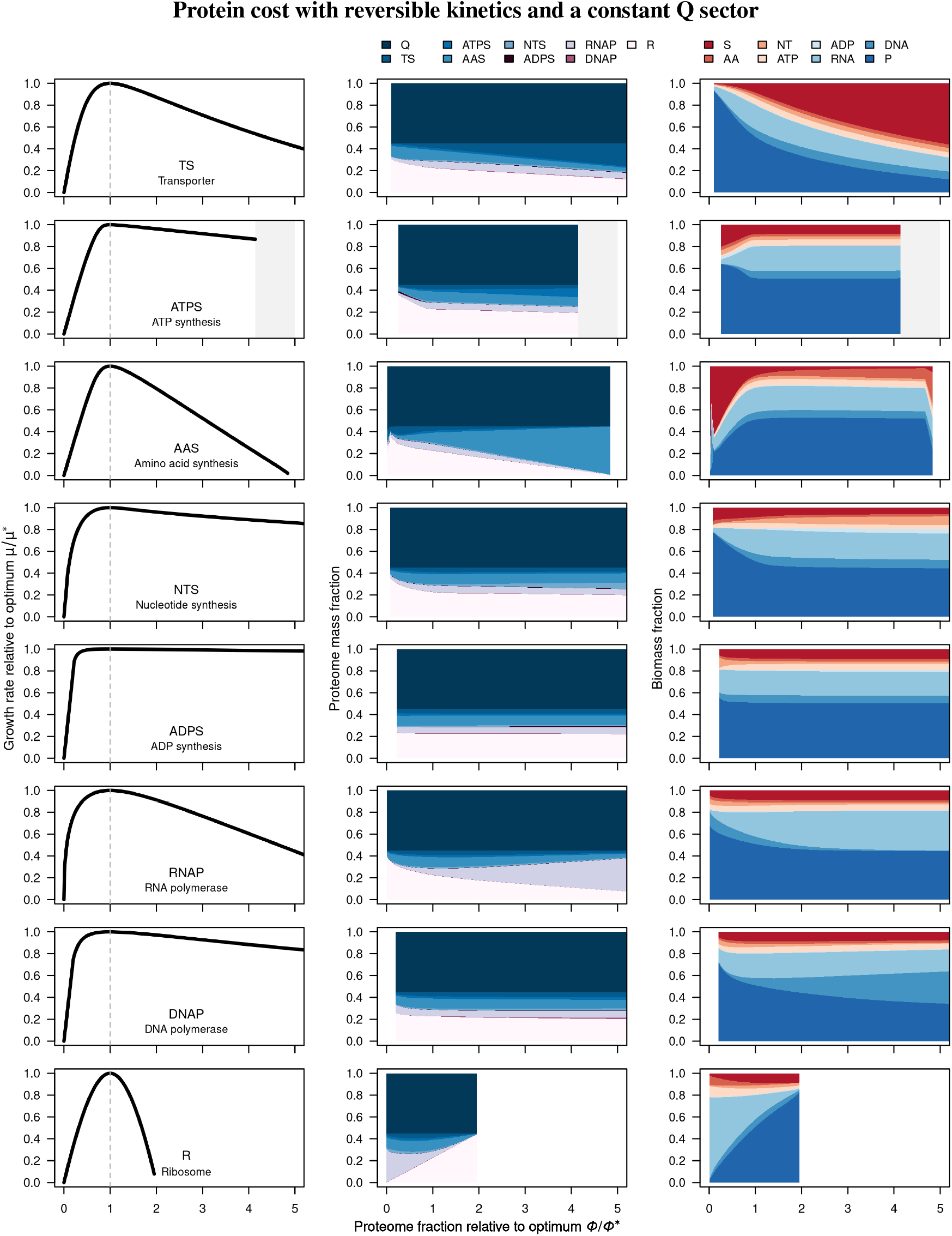
Simulations of protein costs with a model that includes a Q sector (non-growth associated proteins). The Q sector was modeled by adding a column of zeros to the base model and fixed to a constant proteome fraction *ϕ* of 0.55. Kinetic parameters were estimated with the same method as for the base model. Numerical results for ATPS were not found for higher proteome fractions (indicated with the light gray region).

**Fig. S11.**
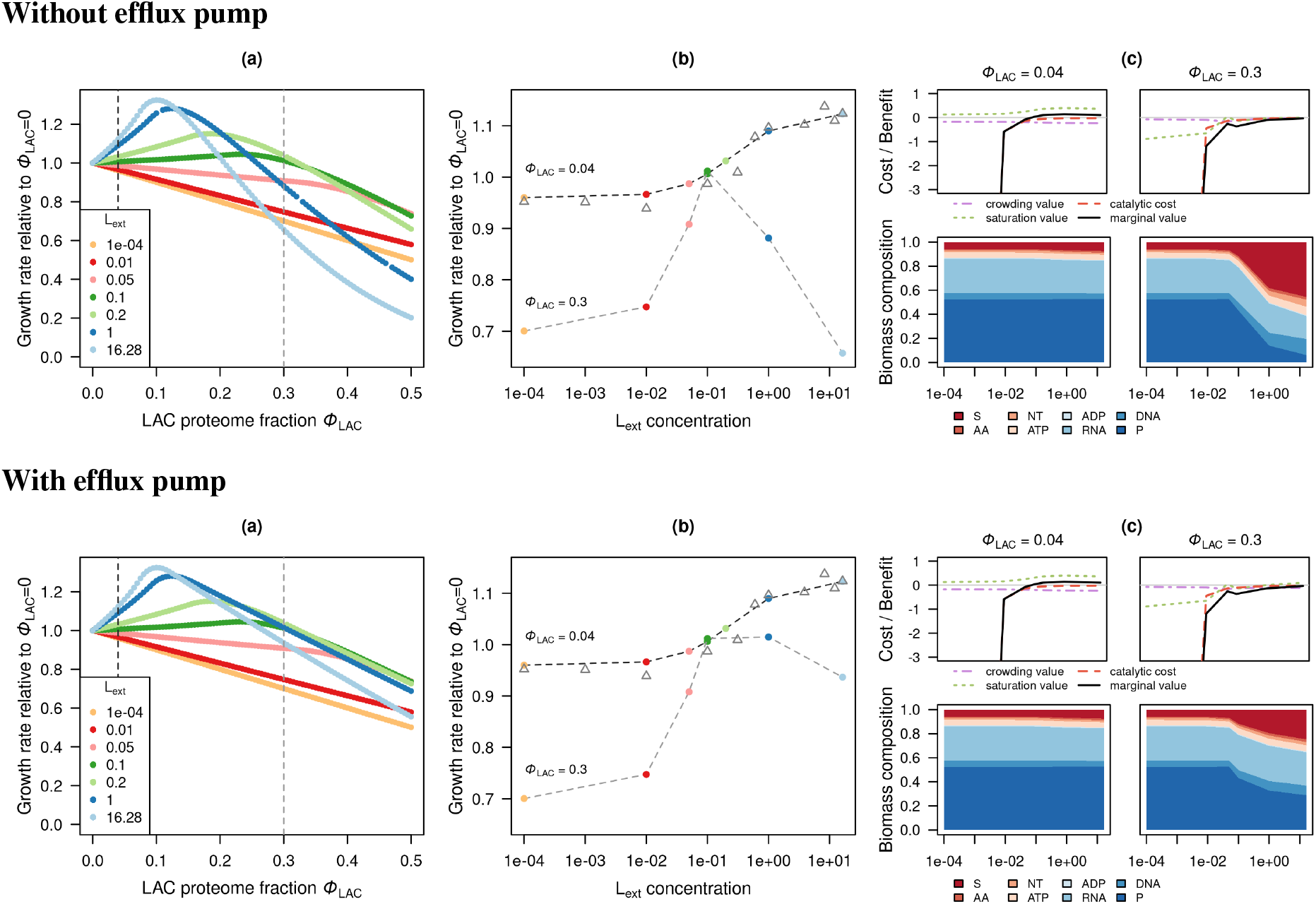
LAC protein growth cost depends on environmental conditions. The two rows show simulations with or without the presence of an efflux pump that exports excess small molecules S out of a cell. In each row **(a)** shows growth rate normalized to *ϕ*_LAC_ = 0 as a function of proteome allocation to LAC (*ϕ*_LAC_) at different L_ext_ concentrations, **(b)** shows the normalized growth rate as a function of L_ext_. The gray triangles are experimental data from (12). **(c) top**: Decomposition of costs and benefits according to Eq. (S1) at fixed proteome fractions (*ϕ* = 0.04, 0.3), showing the crowding value, catalytic cost, saturation value and marginal value. **(c) bottom**: Biomass composition indicates metabolic burden at high *ϕ*_LAC_.

**Fig. S12.**
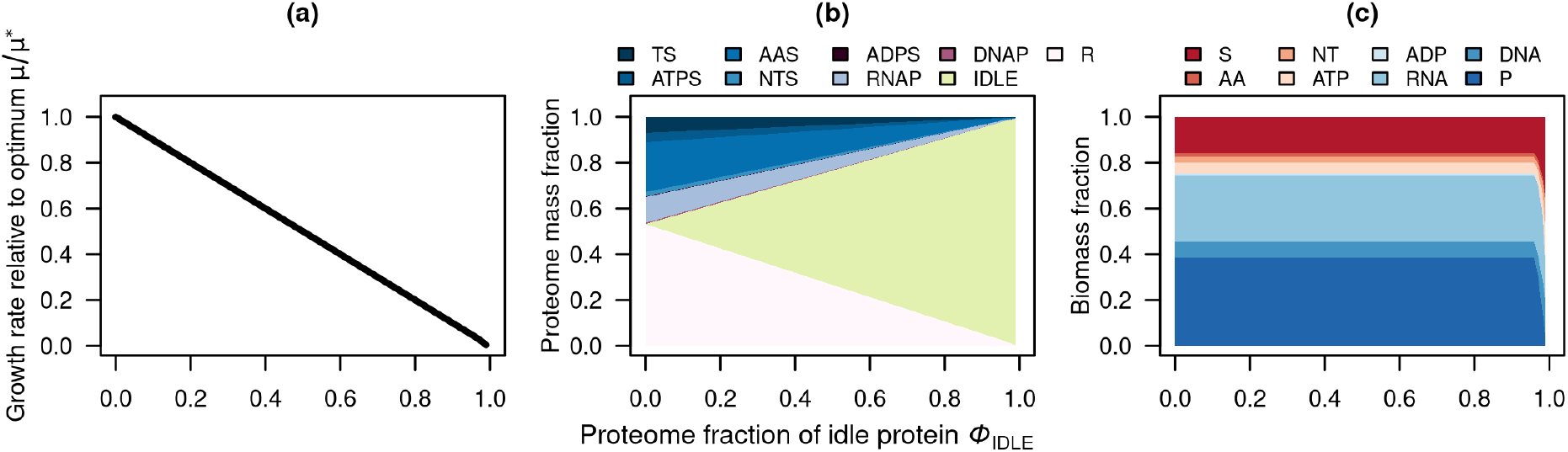
Overexpression of an idle protein leads to an almost linear decrease in growth rate. In contrast to Figure 4, where the non-growth-associated proteome (Q sector) is included, the growth rate decreases to zero as *ϕ* approaches one.

